# Lhx2 is a progenitor-intrinsic modulator of Sonic Hedgehog signaling during early retinal neurogenesis

**DOI:** 10.1101/2021.04.24.441277

**Authors:** Xiaodong Li, Patrick J. Gordon, John A. Gaynes, Alexandra W. Fuller, Randy Ringuette, Clayton P. Santiago, Valerie A. Wallace, Seth Blackshaw, Pulin Li, Edward M. Levine

## Abstract

An important question in organogenesis is how tissue-specific transcription factors interact with signaling pathways. In some cases, transcription factors define the context for how signaling pathways elicit tissue- or cell-specific responses, and in others, they influence signaling through transcriptional regulation of signaling components or accessory factors. We previously showed that during optic vesicle patterning, the Lim-homeodomain transcription factor Lhx2 has a contextual role by linking the Sonic Hedgehog (Shh) pathway to downstream targets without regulating the pathway itself. Here, we show that during early retinal neurogenesis, Lhx2 is a multilevel regulator of Shh signaling. Specifically, Lhx2 acts cell autonomously to control the expression of pathway genes required for efficient activation and maintenance of signaling in retinal progenitor cells. The Shh co-receptors Cdon and Gas1 are candidate direct targets of Lhx2 that mediate pathway activation, whereas Lhx2 directly or indirectly promotes the expression of other pathway components important for activation and sustained signaling. We also provide genetic evidence suggesting that Lhx2 has a contextual role by linking the Shh pathway to downstream targets. Through these interactions, Lhx2 establishes the competence for Shh signaling in retinal progenitors and the context for the pathway to promote early retinal neurogenesis. The temporally distinct interactions between Lhx2 and the Shh pathway in retinal development illustrate how transcription factors and signaling pathways adapt to meet stage-dependent requirements of tissue formation.

## Introduction

The Sonic Hedgehog (Shh) signaling pathway is essential for the patterning, growth, and histogenesis of multiple tissues. Deregulation resulting in hyperactivation drives tumor growth and hypoactivation leads to congenital brain malformations including holoprosencephaly (Hong and Krauss, 2018; Scales and de Sauvage, 2009). The canonical pathway is composed of core and accessory components, which contribute to ligand production, availability, reception, intracellular signaling, and transcriptional regulation of target genes (reviewed in (Briscoe and Thérond, 2013; Kong et al., 2019; Ramsbottom and Pownall, 2016). At its simplest, Shh signaling occurs when secreted ligand binds to its cognate Patched receptor, relieving inhibition of the Frizzled class GPCR transmembrane protein Smoothened (Smo) (Fig. 1A). In turn, an intracellular cascade blocks the proteolytic processing of the GLI zinc-finger transcription factors, Gli2 and Gli3, converting them from transcriptional repressors to activators with Gli3 the predominant repressor and Gli2 the predominant activator (Lipinski et al., 2006). The net result is the expression of downstream genes which includes the third mammalian GLI paralog Gli1, which like Gli2, functions as a transcriptional activator. Gli1 contributes to feedback regulation after signaling is initiated with primary transcriptional targets being itself (positive feedback), or three negative feedback regulators, *Patched 1* (*Ptch1*)*, Patched 2* (*Ptch2*), and *Hedgehog Interacting Protein* (*Hhip*). These feedbacks contribute to steady state signaling (Lai et al., 2004; Li et al., 2018), and allow the expression levels of *Gli1*, *Ptch1*, *Ptch2*, and *Hhip* to be used as readouts of signaling.

**Figure 1:**
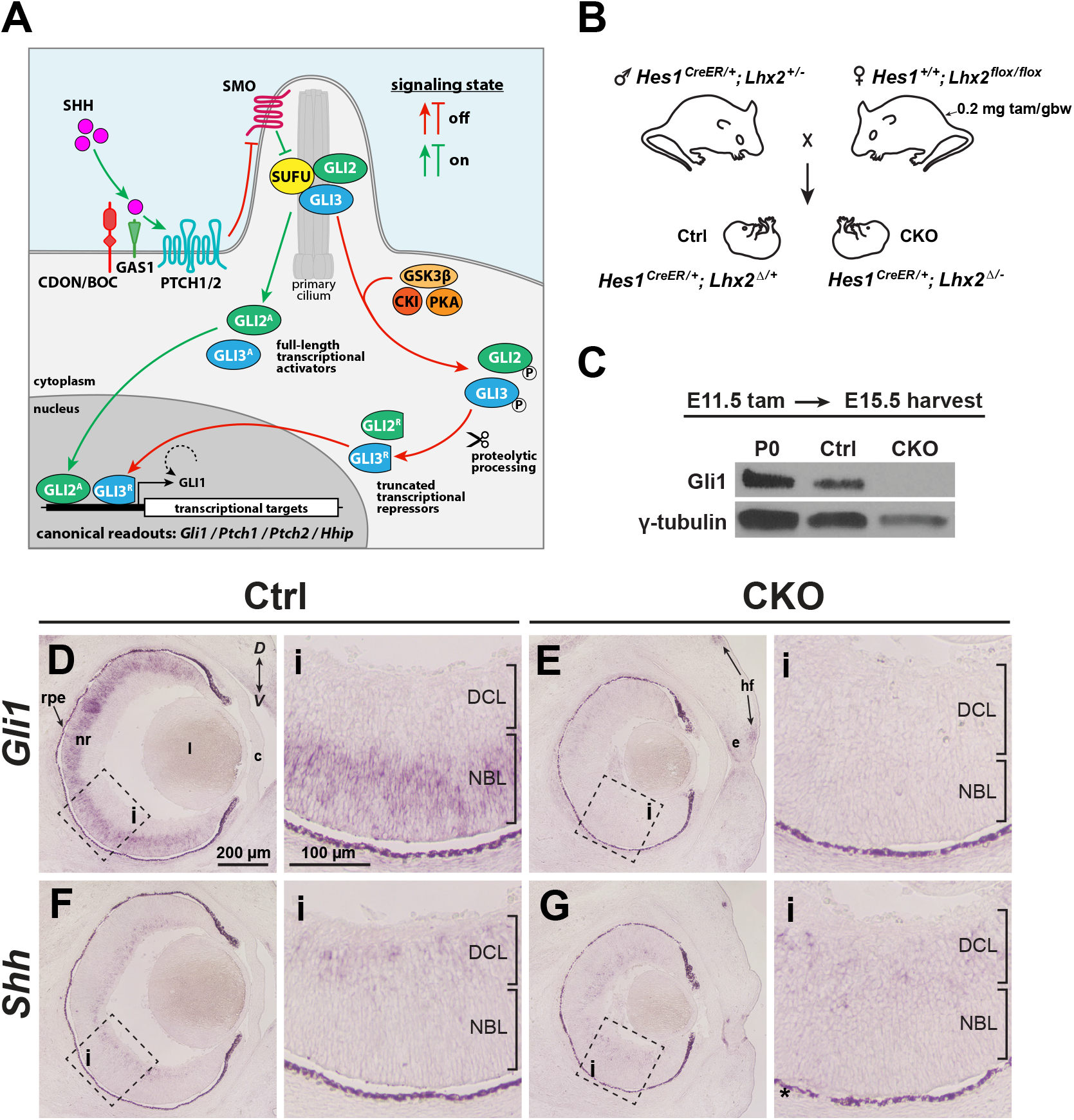
*Lhx2* is required for Gli1 expression in RPCs. **(A)** Overview of Hh signaling. See text for details and Briscoe and Thérond (2013), Ramsbottom and Pownall (2016), and Kong et al. (2019) for more comprehensive pathway illustrations and descriptions. **(B)** Genetics and tamoxifen treatment paradigm. Female breeders are also homozygous for *Rosa^ai14/ai14^* and all embryos are *Rosa^ai14/+^*, allowing rapid screening for recombined embryos with tdTomato expression. Recombined *flox* alleles are indicated by Δ. **(C)** Western blot for Gli1 protein expression in P0 wild type, E15.5 Ctrl and CKO retinas following tamoxifen treatment at E11.5. γ-Tubulin served as an internal loading control. **(D, E)** *in situ* hybridizations for *Gli1* expression in E15.5 Ctrl (**D**) and CKO (**E**) eyes following tamoxifen treatment at E11.5. **(F, G)** *in situ* hybridizations for *Shh* expression in E15.5 Ctrl (**F**) and CKO (**G**) eyes following tamoxifen treatment at E11.5. Dashed boxes reveal locations of close-up images (i). Dark appearance of the rpe is due to natural pigmentation and does not indicate gene expression. **Abbreviations:** *D*, dorsal; *V*, ventral; nr, neural retina; rpe, retinal pigment epithelium; l, lens; c, cornea; e, eyelid; hf, hair follicle DCL, differentiated cell layer; NBL, neuroblast layer

The ability or *competence* of cells to signal is essential for the pathway to exert its effects at the right time, place and magnitude, and many of the core and accessory pathway components contribute to these properties (Kiecker et al., 2016). For a responding cell to be competent, positive transducers must be expressed prior to signaling (e.g. Smo, Gli2), but kept in the ‘off’ state by negative regulators such as Patched, Suppressor of Fused (Sufu), Gli3, and accessory factors such as Protein Kinase A (Pka), Glycogen Synthase Kinase 3 beta (Gsk3b), and Casein Kinase 1 (Ck1). Pathway activation is not achieved, however, by simple ligand binding to Patched, but also requires one of three co-receptors: Cell Adhesion, Oncogene Regulated (Cdon), Brother of Cdon (Boc), or Growth Arrest Specific 1 (Gas1). Thus, pathway regulation is complex even before signaling is initiated. Since many pathway genes are not expressed ubiquitously, understanding how they are regulated in specific contexts can reveal how signalling is tailored to meet the demands of developing tissues and provide insights into developmental timing mechanisms.

In early eye development, Shh signaling is initially required for the regionalization and ventral patterning of the optic neuroepithelium including in the nascent neural retinal domain (Chiang et al., 1996; Gallardo and Bovolenta, 2018; Hernández-Bejarano et al., 2015; Take-uchi et al., 2003; Wang et al., 2015; Zhao et al., 2010). In the newly formed retina, a second interval of signaling occurs in retinal progenitor cells (R<u>P</u>Cs) at the onset of neurogenesis, propagating as a central-to-peripheral wave that is coupled to retinal ganglion cell (R<u>G</u>C) production and RGC-derived *Shh* expression (reviewed in (Wallace, 2008). This coupling is maintained even when RGC production is delayed (Sigulinsky et al., 2008) indicating that ligand availability sets the developmental timing of pathway activation because RPCs are competent to signal prior to ligand exposure. How this competence is established in RPCs at the start of retinal neurogenesis has not been addressed.

The LIM-homeodomain transcription factor *Lim-homeobox 2* (*Lhx2*) is a multifunctional regulator of retinal development. Initially expressed in the eye field, Lhx2 expression persists throughout retinal development in RPCs, becoming restricted to Muller glia and a subset of amacrine cells (de Melo et al., 2012; Gordon et al., 2013; Tétreault et al., 2009; Viczian et al., 2006; Yun et al., 2009). Initially, Lhx2 is required for optic vesicle patterning and regionalization, lens specification, and optic cup morphogenesis (Hägglund et al., 2011; Porter et al., 1997; Roy et al., 2013; Seth et al., 2006; Tétreault et al., 2009; Yun et al., 2009; Zuber et al., 2003). Lhx2 directs these processes through cell autonomous and nonautonomous mechanisms, in part through regulation of optic vesicle-derived expression of the Bone Morphogenetic Proteins BMP4 and BMP7 (Yun et al., 2009). After optic cup formation, Lhx2 nonautonomously directs lens development in part through regulation of retinal-derived expression of FGFs, and possibly BMP4 (Thein et al., 2016). Toward the end of retinal histogenesis, Lhx2 is required at multiple steps in the formation of Muller glia, the sole RPC-derived glial cell type (de Melo et al., 2016a; de Melo et al., 2016b). In this case, an interaction with Notch signaling is partly responsible, through Lhx2-dependent expression of ligand (*Notch1*), receptors (*Dll1*, *Dll3)*, and downstream transcriptional effectors (*Hes1*, *Hes5*) (de Melo et al., 2016b). Thus, Lhx2’s multifunctional nature is defined in part by its interactions with developmental signaling pathways.

We previously reported that RPC-directed *Lhx2* inactivation during embryonic stages of retinal neurogenesis in mice disrupted histogenesis through reduced proliferation, a failure to maintain the progenitor pool, and altered fated precursor cell production (Gordon et al., 2013). Most notably, inactivation of Lhx2 at the onset of retinal neurogenesis caused the overproduction of RGCs. These phenotypic features are similar to what occurs when Shh signaling is inhibited at the same developmental stage in the mouse retina or comparable stage in the chick retina (Wang et al., 2005; Zhang and Yang, 2001), which prompted us to investigate a potential connection between Lhx2 and Shh signaling.

## Results

### Lhx2 is required for Gli1, but not Shh, expression at the start of retinal neurogenesis

Gli1 is functionally dispensable in mice, but its expression depends on Shh signaling in many tissues including the retina (Bai et al., 2002; Furimsky and Wallace, 2006; Marigo et al., 1996; McNeill et al., 2012; Park et al., 2000; Sigulinsky et al., 2008; Wang et al., 2002). Because of its well-established role as a readout of pathway activity, we first asked if Gli1 expression was altered by the loss of Lhx2 function. *Lhx2* was temporally inactivated in RPCs using the *Hes1^CreER^* driver by administering tamoxifen to timed pregnant dams at E11.5 (Fig. 1B), coincident with the onset of retinal neurogenesis and activation of Shh signaling in RPCs. At E15.5, Gli1 protein and mRNA expression levels were markedly reduced in the CKO (Fig. 1C-E). Tamoxifen treatment at E12.5 showed a similar reduction in *Gli1* expression although sparse expression was observed dorsally (Supplemental Fig. 1). Similar to the control (Ctrl) retina, *Shh* expression was detected across the CKO retina (Fig. 1F,G), indicating that the drop in *Gli1* was not due to a lack of *Shh* expression, although this does not rule out problems with ligand availability. We therefore set out to determine if the drop in *Gli1* was due to a role for Lhx2 in promoting Shh signaling, and if so, at what level.

### Lhx2 inactivation alters the expression of multiple Shh pathway components

Bulk RNA sequencing (RNA-seq) was done on retinas isolated from E15.5 embryos following tamoxifen treatment at E11.5 (Fig. 2A). Approximately 12,300 features (i.e. protein coding genes, pseudogenes, lncRNAs) were identified and examined for differential expression using DESeq2 (Fig. 2B; Supplemental Table 1). Overall, 2210 differentially expressed features remained after applying a false discovery rate (FDR) cutoff of 0.001, with 2161 identified as protein coding genes, the next largest category being lncRNAs at 20. Of these 2210 features, collectively referred to as differentially expressed genes (DEGs), 1184 were downregulated in the CKO within the log2 transformed fold change (log2FC) interval from -5.69 to -0.19, and 1026 DEGs were upregulated within the log2FC interval from 0.17 to 7.69. These statistics are in line with Lhx2 being an essential developmental transcription factor.

**Figure 2:**
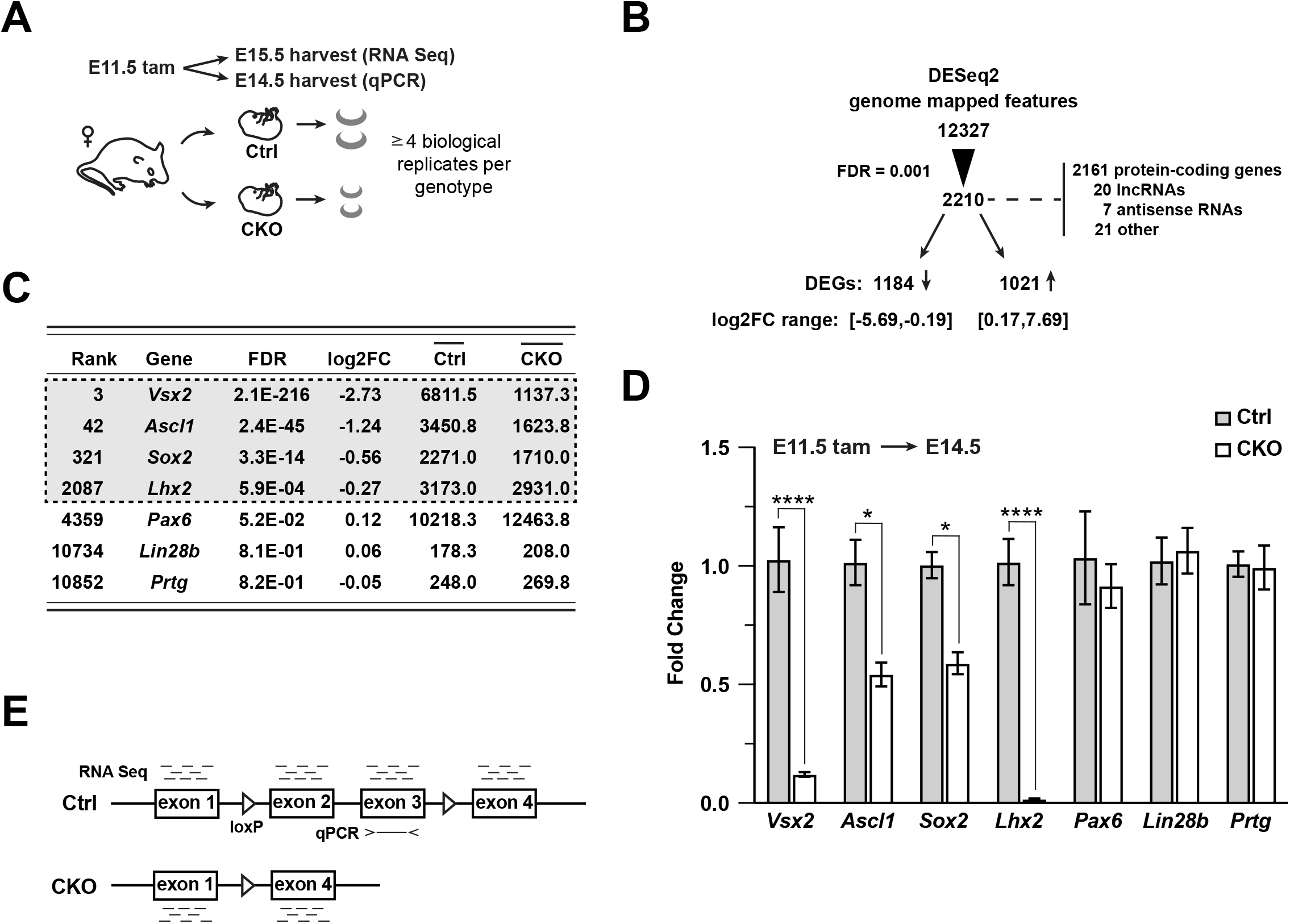
Gene expression changes due to *Lhx2* inactivation. **(A)** Schematic of experimental design for RNA sequencing and qPCR. **(B)** Summary of DESeq2 analysis of RNA sequencing datasets. **(C)** DESeq2-derived statistics for progenitor genes with requirements during early retinal neurogenesis. Genes in gray box were within the 0.001 FDR cutoff for differential expression. Averaged counts per gene are shown in the last two columns. **(D)** Relative expression for progenitor genes at E14.5 by qPCR as a function of the fold change from the mean of control for each gene. Only significant comparisons are noted (*, p_adj_ < 0.05; ****, p_adj_ < 0.0001; see Supplemental Table 3 for statistics). **(E)** Schematic showing the differences in the relative expression of *Lhx2* by RNA sequencing and qPCR. The qPCR primers are located in exon 3, which is deleted by Cre recombination, making the mutant transcript undetectable by qPCR. The mutant transcript is expressed and detected by RNA sequencing.

Semi-quantitative RT-PCR (qPCR) was used to validate the RNA-seq data. In addition to *Lhx2*, the RPC expressed genes *Vsx2*, *Ascl1*, and *Sox2* were downregulated whereas *Pax6*, *Lin28b* and *Prtg* remain unchanged (Fig. 2C). Compared to the RNA-seq data, all genes showed similar trends in relative expression except for *Lhx2*, which showed a more pronounced reduction by qPCR (Fig. 2D; Supplemental Table 3). Because the primers are located in third exon, which is deleted in the CKO allele, the discrepancy between the RNA-Seq and qPCR is consistent with persistent expression of the mutant transcript rather than incomplete recombination (Fig. 2E). Overall, the qPCR data aligns well with the RNA-Seq data and the applied FDR cutoff serves as a reliable indicator of differential expression.

We analyzed the top 2210 DEGs using a pathway topology algorithm based on curated KEGG pathways (Canonical Pathways, IPA, Qiagen). Applying post-algorithm cutoffs for adjusted p-values ≤0.05 (-log(padj) ≥1.3) and absolute Z-scores ≥2.5 (Fishers exact test), 12 pathways were identified with predicted changes in activity (Fig. 3A). In general, activated pathways were associated with neuronal differentiation (orange bars) and inhibited pathways with cell cycle progression (blue bars), consistent with the CKO phenotype (Gordon et al., 2013). Shh signaling was identified as an inhibited pathway (Fig. 3A) with approximately 35% of the genes identified as DEGs (ratio curve, right Y-axis). We also examined an Lhx2 ChIP-Seq dataset from FACS-enriched E14.5 wild type RPCs with clusterProfiler to identify candidate KEGG pathways associated with Lhx2 binding based on assigning Lhx2 peaks to the nearest gene locus (see below for criteria used for peak to gene association) (Yu et al., 2012; Zibetti et al., 2019). The Shh pathway (listed Hedgehog signaling) was scored as a significantly associated pathway and was the only pathway to be present on both lists with the employed criteria (Fig. 3B; Supplemental Table 2). The *Wnt/β -catenin* and *axon guidance* pathways had smaller p-values than Shh signaling in IPA and clusterProfiler, but their Z-scores for predicted activity states failed to reach the cutoff (Supplemental Table 2). These observations support a link between Lhx2 and the Shh pathway at the level of gene expression, chromatin regulation and signaling activity.

**Figure 3:**
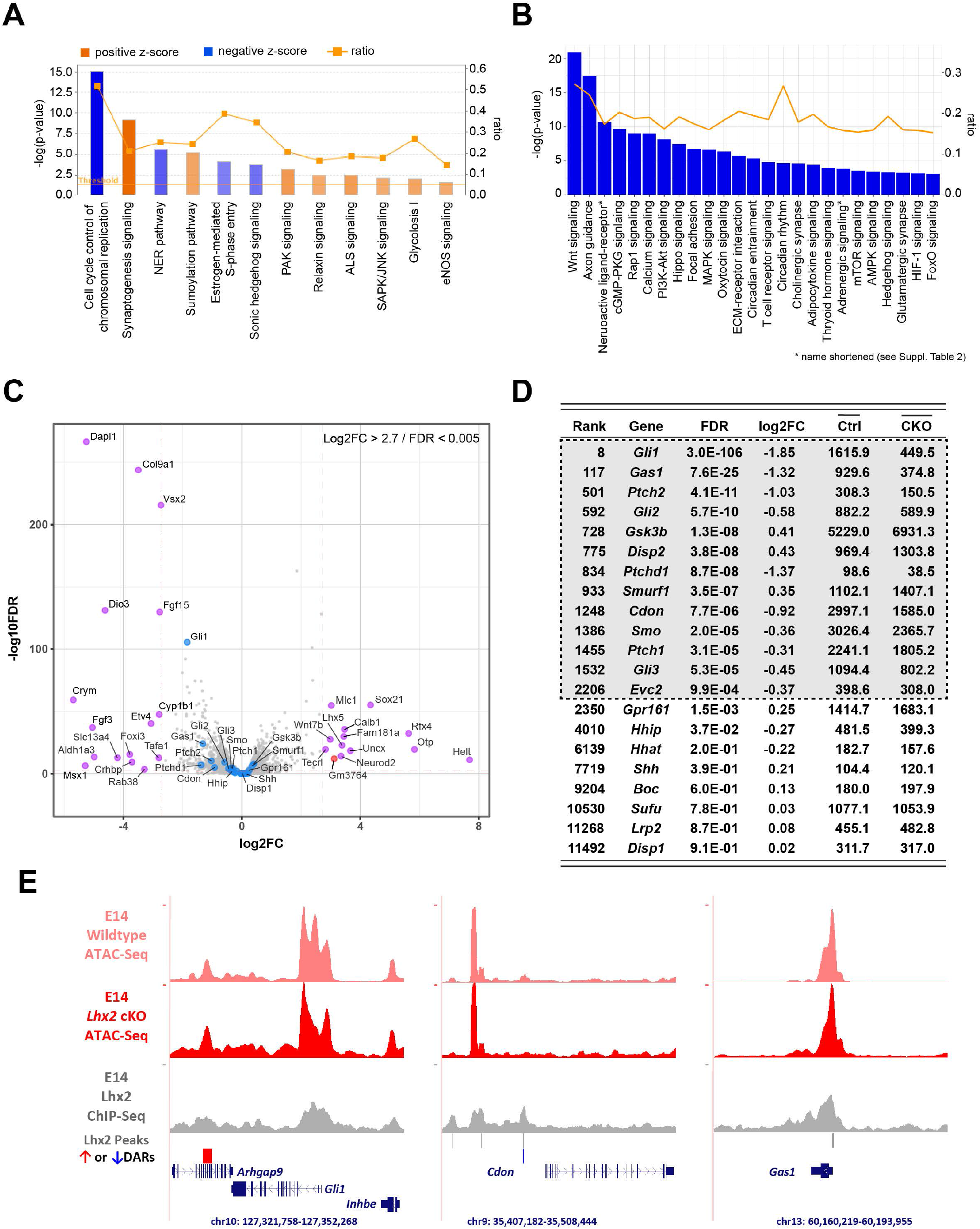
Lhx2 is required for the expression of multiple Hedgehog pathway genes. **(A)** Canonical KEGG pathways analysis for DEGs with the 0.001 FDR cutoff. Pathways shown are those with a -log(P_adj_) ≥ 1.3 <u>and</u> an absolute z-score ≥ 2.5 to show only top-ranked pathways predicted to be inhibited (blue bars) or activated (orange bars) with higher color intensity (saturation) directly correlated with z-score. Connected line indicates the ratio of DEGs represented in each KEGG pathway pathway list (right y-axis). **(B)** KEGG pathways associated with Lhx2 ChIPSeq peak distribution from E14.5 mouse RPCs. The connected line indicates the ratio of Lhx2-associated genes represented in each pathway (right y-axis). **(C)** Volcano plot showing selected Hh pathway genes (blue dots) relative to other DEGs. The most divergent DEGs gated on FDR (0.005) <u>and</u> absolute log2FC (2.7) cutoffs are highlighted (purple indicates mRNAs; red dot indicates lncRNA). **(D)** DESeq2-derived statistics for Hh pathway genes. Genes in gray box were within the 0.001 FDR cutoff. Averaged counts per gene are shown in the last two columns. **(E)** ATAC-Seq and Lhx2 ChIP-Seq tracks at *Gli1*, *Cdon*, and *Gas1* loci from E14.5 RPCs. Differentially accessible chromatin regions (DARs) in *Lhx2* CKO RPCs compared to control are indicated by red (open) and blue bars (closed). Lhx2 ChIPSeq peak calls are indicated by gray bars.

To assess this potential link further, we compiled a list of genes with established and direct roles in Shh signaling. None of the genes were among the most divergent in the RNA-Seq dataset using a conservative fold change cutoff of log2FC >2.7 for visualization purposes (Fig. 3C). However, *Gli1* was ranked 8^th^ overall by DESeq2 and was the top-ranked DEG in the Shh pathway (Fig. 3D). Multiple pathway genes in the list qualified as DEGs and have established functions specifically in responding cells (Fig. 3D, gray box). At the receptor level, *Cdon*, *Gas1, Ptch1*, *Ptch2,* and *Smo* were reduced in the CKO. Lower expression of *Gas1, Cdon,* and *Smo* could negatively impact signaling, which is consistent with the overall reduction in Shh signaling in the CKO retina. On the other hand, reduced *Ptch1* and *Ptch2* should promote signaling, but since this was not the case, the decreases in *Ptch1* and *Ptch2* expression are likely due to reduced signaling, consistent with their expression levels serving as readouts of pathway activity.

At the intracellular level of the pathway, *Gli2* expression was reduced in the CKO, but *Gli3*, a likely direct target of *Lhx2* (Zibetti et al., 2019) was also reduced, potentially offsetting the drop in *Gli2* (Furimsky and Wallace, 2006). *Gli2* expression is not typically considered to be dependent on Shh signaling, but its expression level declined following RPC-specific *Smo* inactivation (Sakagami et al., 2009), leaving open the possibility that it is transcriptionally regulated by Shh signaling in RPCs. Of the two upregulated DEGs, only *Gsk3b* is predicted to inhibit signaling by promoting the processing of Gli3 into the repressor isoform. Its inhibitory function, however, is normally overridden by ligand-based activation (Briscoe and Thérond, 2013). Thus, *Gsk3b* activity could contribute to reduced signaling in the CKO, but only if the pathway is disrupted upstream.

These observations suggest that expression of *Gas1, Cdon, Smo, Gli3,* and *Gsk3b* are preferentially dependent on Lhx2, and *Ptch1* and *Ptch2* preferentially dependent on signaling via Gli1 and Gli2. The reduced levels of *Gli1* and *Gli2* could be independent consequences of *Lhx2* inactivation, reduced Shh signaling, or a combination of the two. Since Lhx2 is a transcription factor with pioneer activity, we assessed Lhx2 chromatin binding using an Lhx2 ChIP-Seq dataset from E14.5 wild type RPCs, and analyzed changes in chromatin accessibility due to *Lhx2* inactivation with ATAC-Seq datasets from E14.5 control and *Lhx2* CKO RPCs (Zibetti et al., 2019). We employed two criteria to associate genes with ChIP-Seq peaks and ATAC-Seq derived differentially accessible chromatin regions (DARs): (i) peaks had to be located within the gene body or within 10 kb upstream or downstream, or (ii) the assigned gene was the closest gene to the peak. By both criteria, ChIP-Seq peaks were not associated with *Gli1, Gli2*, *Gsk3b, Ptch2*, or *Smo.* DARs, however, were associated with *Gli1*, *Gli2,* and *Gsk3b* suggesting indirect regulation by Lhx2 (Fig 3E; Supplemental Fig. 2). In contrast, Lhx2 binding was associated with *Cdon*, *Gas1, Ptch1,* and *Gli3* (Fig 3E; Supplemental Fig. 2), and several ChIP-Seq peaks aligned with DARs, suggesting direct regulation by Lhx2 (Fig 3E; Supplemental Fig. 2) (Zibetti et al., 2019). ChIP-Seq peaks were also associated with Shh pathway genes with low DESeq2 ranking or weak expression (Supplemental Figs. 2 and 3). In sum, the expression of multiple Shh pathway genes is altered following *Lhx2* inactivation, with several likely to be directly dependent on Lhx2 and others more generally regulated by Shh signaling.

### Lhx2 does not regulate Shh ligand availability

Shh signaling in the retina is an example of *intra*-lineage signaling, where the ligand producing cells (RGCs) are the direct descendants of the responding cells (RPCs). This configuration, in effect, places Shh upstream and downstream of the pathway (Fig. 4A). Although RGCs are overproduced in the CKO and the expression of genes involved in Shh production did not reach the cutoff for confident DEG designation (Fig. 3D), it stands that mRNA expression levels are insufficient for predicting ligand bioavailability, especially given the importance of posttranslational mechanisms in ligand modification, secretion, and presentation (Briscoe and Thérond, 2013). Furthermore, *Lhx2* loss of function could cause cryptic changes in extracellular factors that could negatively impact signaling. A more direct approach is to functionally test the endogenous Shh ligand with minimal disruption to the extracellular environment. To do this, we adapted a biosynthetic system engineered to model gradient formation and signaling dynamics in Shh-responsive NIH3T3 cells (Li et al., 2018). In this reporter system, *Ptch1* is expressed under the control of a stably integrated doxycycline-regulated expression cassette in a *Ptch1* mutant background and signaling is reported by the expression of H2B-mCitrine under the control of 8 tandem GLI binding sites (Fig. 4B). Referred to as an *open-loop* circuit, this configuration eliminates Ptch1-mediated negative feedback and expands the dynamic range of mCitrine reporter activity in these cells and are referred to as *open-loop cells*) (Li et al., 2018). This system has the potential to directly identify deficiencies in the bioavailability of endogenous ligand.

**Figure 4:**
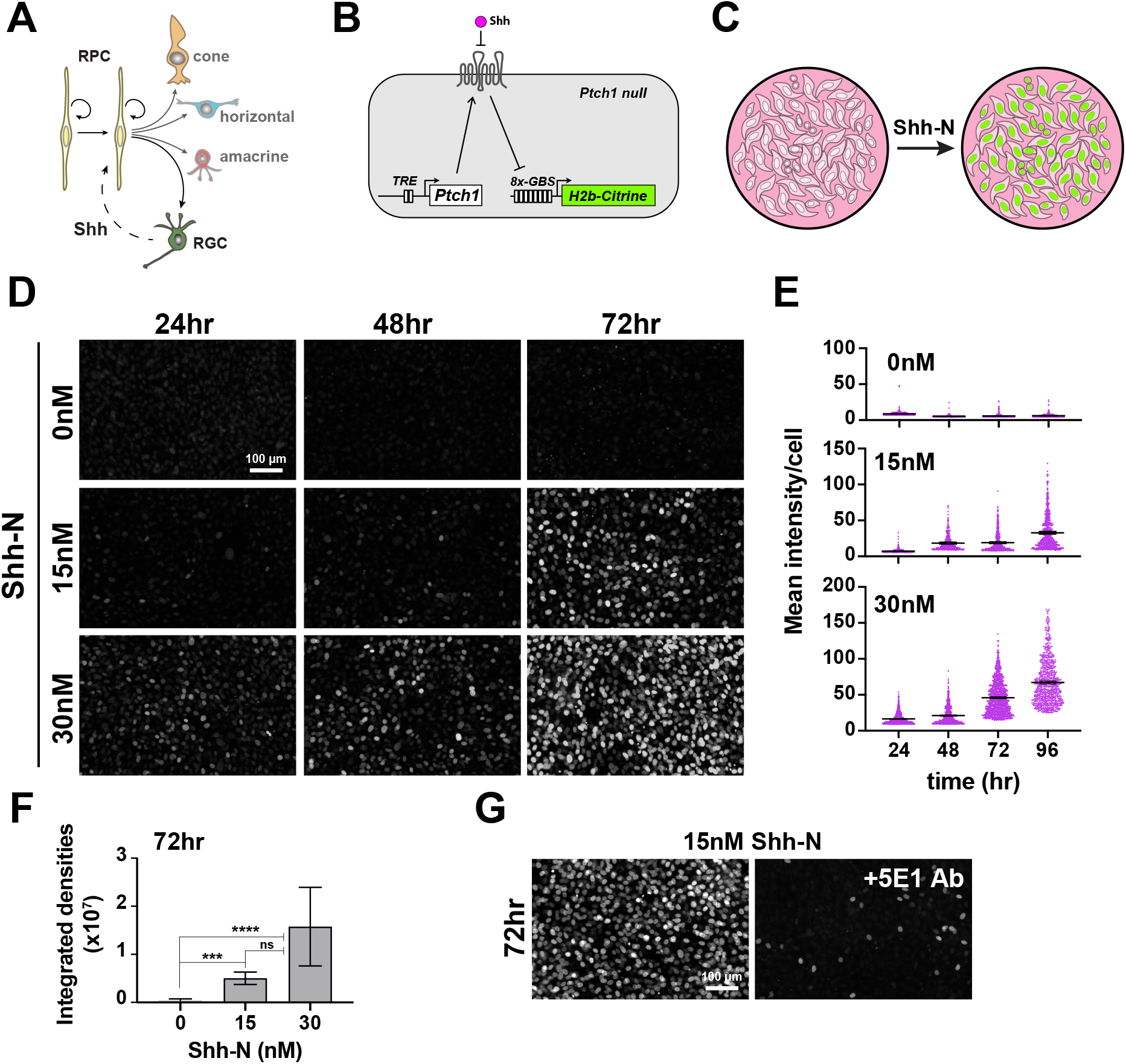
Evaluation of a live cell reporter system to test ligand availability. **(A)** *Intra-*lineage architecture of Hh signaling at the start of retinal neurogenesis. RPCs initiate neurogenesis and begin generating RGCs, the Shh producing cells. RPCs are the responder cells, placing ligand upstream and downstream of RPCs. **(B)** Configuration of the open loop circuit in the responder NIH3T3 *Ptch1-*null cell line. Ptch1 is produced from a doxycycline regulated transgene. Hh ligand binds Ptch1, activating intrinsic signaling as well as Gli-dependent expression of mCitrine fused to Histone H2b through 8-multimerized Gli1 binding sites. **(C)** Addition of recombinant Shh-N to a confluent monolayer of responder cells activates nuclear mCitrine expression. **(D)** Dose response and time course for accumulation of mCitrine in open loop responder cells. Fields were randomly selected from the same cultures for each timepoint. **(E)** Plots showing the accumulation of mCitrine+ cells over time for each dose tested as a function of average mean fluorescence intensity. Means are shown for each condition (lines). **(F)** Dose response at 72 hr as a function of the sum of the integrated densities of the mCitrine+ cells. (ns, not significant; ***, p_adj_ < 0.001; ****, p_adj_ < 0.0001; see Supplemental Table 4 for statistics) **(G)** Addition of the Shh ligand-blocking monoclonal antibody, 5E1, diminishes mCitrine expression.

To assess the suitability of this reporter system, open-loop cells were grown to confluence followed by addition of recombinant human Shh (N-terminal fragment, C24II variant; referred to as Shh-N) and monitored for mCitrine fluorescence (Fig 4C). We observed dose-dependent increases in mCitrine expression at 15nM and 30nM Shh-N over the course of 96hr with 72hr being a suitable time point to detect robust expression (Fig. 4D-F; Supplemental Table 4). Addition of the Shh-blocking IgG antibody, 5E1, to the culture medium suppressed mCitrine expression (Fig. 4G), demonstrating a continued requirement for ligand in the open-loop cells even at the lowest levels of Ptch1 expression (without doxycycline treatment).

We next tested the ability of retinal tissue to stimulate signaling in open-loop cells. Prior work showed that mCitrine+ cells extended several cell diameters away from the cellular source of ligand as long as cell contact was uninterrupted (Li et al., 2018). We therefore developed a coculture paradigm in which whole retinal tissue was flat mounted onto the underside of a PTFE transwell insert, placing the retina in direct contact with the confluent monolayer of open-loop cells (Fig. 5A). We initially tested E18.5 wild type retina because RGCs are abundant and Shh signaling extends across the retina by this age (Sigulinsky et al., 2008) (Supplemental Fig. 4). Open-loop cells responded well to retinal Shh ligand with more mCitrine+ cells observed when the basal surface of the retina was placed in direct apposition, consistent with the location of RGCs. Thus, this reporter system can be used to assess ligand activity in the intact retina especially when the open-loop cells are in close proximity to RGCs, the Shh-producing cells.

**Figure 5:**
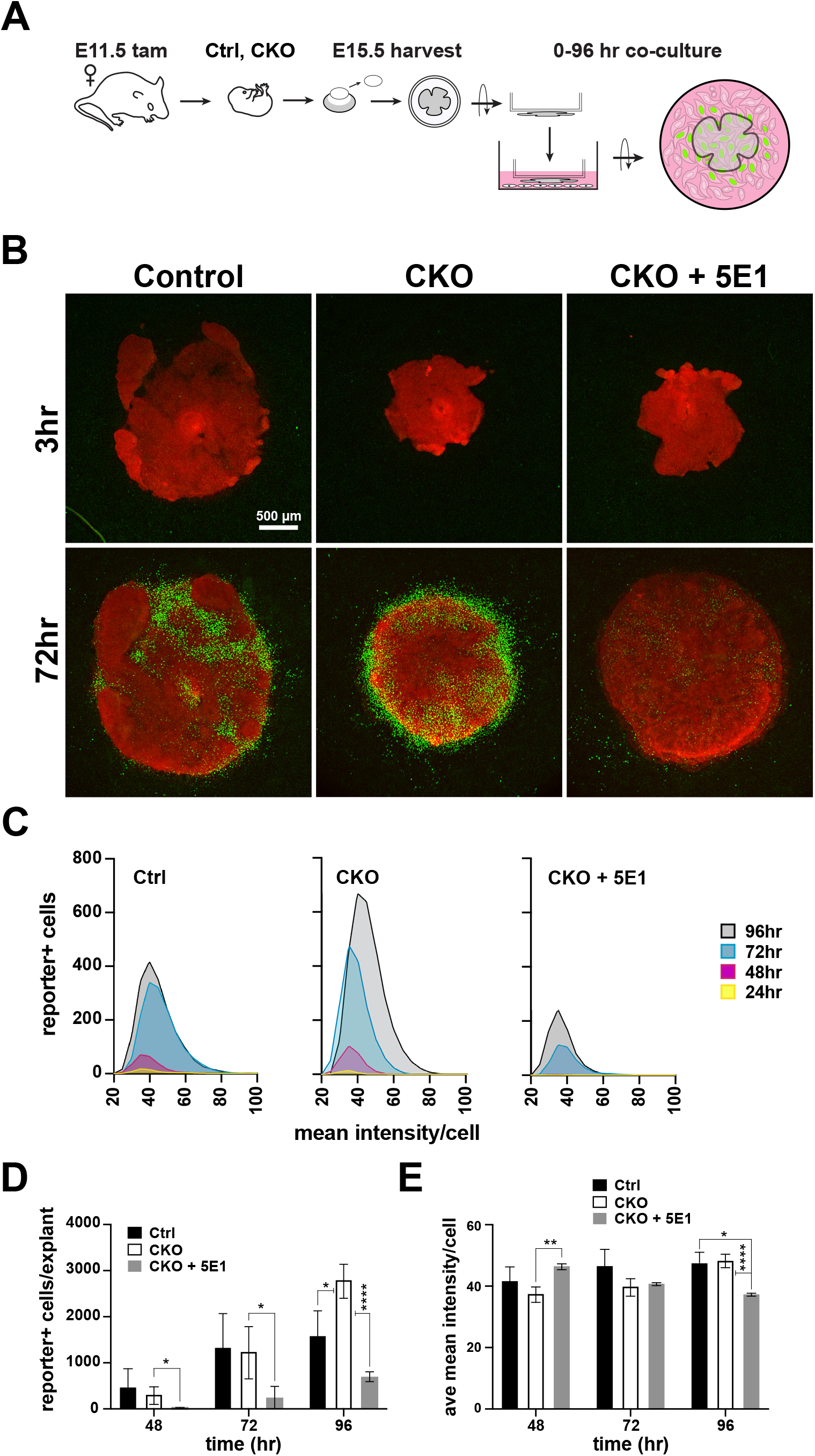
Endogenous Shh activity is robust following *Lhx2* inactivation. **(A)** Experimental design for coculture of retinal explants and open loop responder cells. Freshly dissected embryonic retina with lens removed is flat mounted to the underside of a transwell insert and with the apical or basal surface of the retina placed into direct contact with a confluent monolayer of NIH3T3 open loop responder cells. mCitrine expression accumulates in responders that receive Shh from the retina. Pilot experiments revealed a more robust response when RGCs (basal surface) were directly apposed to responder cells (Supplementary Fig. 4). **(B)** Cocultures for control (left), CKO (middle), and CKO incubated with 5E1 antibody (right) at 3 and 72hr. Retinal tissues are tdTomato positive (red) and mCitrine positive nuclei are green. **(C)** Lowess-smoothed histograms showing the accumulation and fluorescence intensity distributions of mCitrine+ responder cells at each timepoint during the co-culture period. The histograms are for the cocultures shown in A. **(D)** Quantification of mCitrine+ cells at 48, 72, and 96 hr. Comparisons were done within timepoints only and the significant differences are shown (*, p_adj_ < 0.05; **, p_adj_ < 0.01; ****, p_adj_ < 0.0001; see Supplemental Table 4 for statistics). **(E)** Quantification of the average mCitrine fluorescence intensities per cell at 48, 72, and 96 hr. Comparisons were done within timepoints only and the significant differences are shown (*, p_adj_ < 0.05; **, p_adj_ < 0.01; ****, p_adj_ < 0.0001; see Supplemental Table 4 for statistics).

If reduced signaling was not due to impaired ligand activity or availability, then open-loop cells should respond similarly to ligand from CKO retinas compared to control. This response could be reflected in the number of mCitrine+ cells, the expression levels of mCitrine, or in the kinetics of mCitrine accumulation. Tamoxifen was administered at E11.5 and retinas were harvested at E15.5 and placed onto confluent open-loop cell monolayers and longitudinally imaged at 24 hr intervals by widefield epifluorescence (Fig 5A). mCitrine expression was not observed at the start of the culture period but was readily apparent in control and CKO cocultures by 72hr (Fig. 5B) and in close proximity to all explants tested (10 control, 13 CKO). mCitrine expression was ligand dependent, revealed by the reduction in mCitrine+ cells when the 5E1 antibody was added to the culture medium (Fig. 5B). Using representative explants for each condition, the accumulation and fluorescence intensities of mCitrine+ cells were plotted over time (Fig. 5C). In general, the behavior of the open-loop cells exhibited similar temporal characteristics but CKO explants supported the highest number of mCitrine+ cells at 96hr, and the accumulation of mCitrine+ cells in the presence of 5E1 was notably reduced at all times (Fig. 5C). To quantify these observations, the total number of mCitrine+ cells per explant and the mean fluorescence intensity per cell were assessed with aggregated data from multiple cocultures (Fig. 5D,E; Supplemental Table 4). Ctrl and CKO explants promoted similar numbers of mCitrine+ cells at 48hr and 72hr, but more mCitrine+ cells were associated with the CKO explants at 96hr. The number of mCitrine+ cells were significantly reduced in the 5E1-treated CKO cocultures at all three timepoints (Fig. 5D). The fluorescence intensities per cell were generally similar across conditions indicating similar levels of signaling were achieved regardless of the explant’s genotype. Subtle but significant differences in mean fluorescence intensities were observed in the 5E1-treated CKO cocultures (Fig. 5E) but given the low numbers of mCitrine+ cells in this condition (Fig. 5D), the differences could be due to stochastic variation in Ptch1 expression in a small cohort of open-loop cells (Li et al., 2018). In sum, these data reveal that ligand availability is not compromised in the CKO retina and point to a cell-autonomous role for Lhx2 in promoting Shh signaling during early retinal neurogenesis.

### Modulating Smoothened activity ex-vivo stimulates Shh signaling in Lhx2-deficient RPCs

We next asked if the reduced Shh signaling activity in the CKO retina was due to defective intracellular signal transduction. Smo is an obligate component of Shh signaling and functions at the interface of the extracellular and intracellular portions of the pathway (Fig. 1A) (Briscoe and Thérond, 2013). We reasoned that if the pathway was not functional at the level of Smo or downstream, attempts to stimulate signaling via Smo would fail. To address this, we utilized an *ex vivo* organotypic culture paradigm previously used to assess Shh signaling in postnatal day 0 (P0) RPCs (Sigulinsky et al., 2008). Here, explant cultures consisting of whole retina with the lens still attached were established from E14.5 control and CKO embryos and treated with purmorphamine, a small molecule agonist of Smo that bypasses Patched-mediated inhibition (Stanton and Peng, 2010) (Fig. 6A). To minimize the potential impact of the tissue phenotype on gene expression levels, the interval between tamoxifen treatment and the start of the culture was shortened to 3 days, which was still sufficient to observe reduced *Gli1* and *Smo* expression in the CKO (Fig. 6B; Supplemental Table 3).

**Figure 6:**
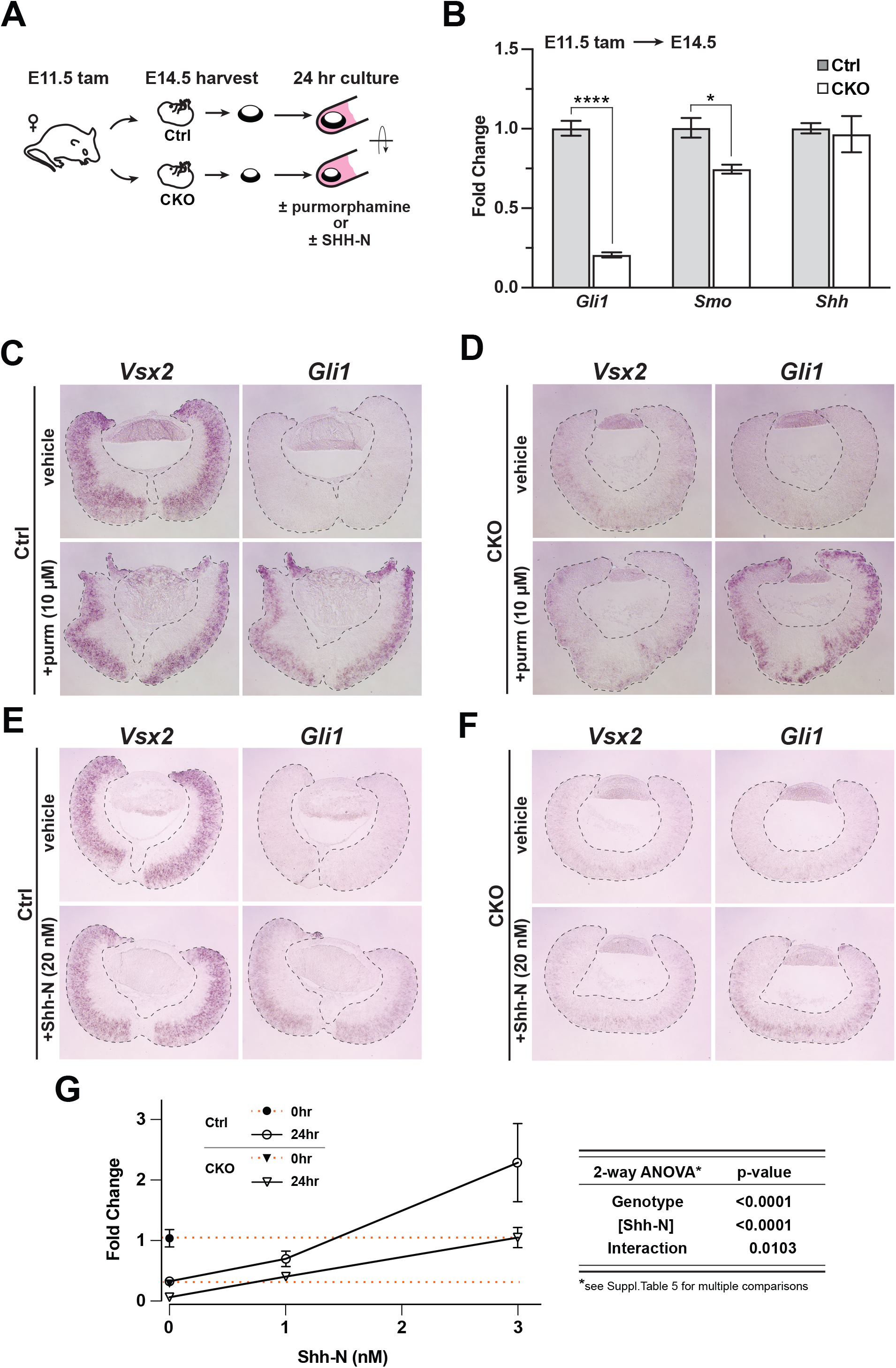
Purmorphamine and Shh-N stimulate Hh signaling in *Lhx2* CKO retinal explants. **(A)** Experimental design of *ex vivo* retina-lens explant cultures. **(B)** Relative expression of *Gli1*, *Smo*, and *Shh* in E14.5 retina by qPCR as a function of the fold change from the mean of control for each gene. Significant comparisons are shown (*, p_adj_ < 0.05; ****, p_adj_ < 0.0001; see Supplemental Table 5 for statistics). **(C-F)** *in situ* hybridizations of *Vsx2* and *Gli1* expression after 24hr in culture to test *Smo* agonist purmorphamine or recombinant Shh-N ligand. **(C)** *Gli1* expression declines in the absence of purmorphamine (vehicle) whereas *Vsx2* is maintained. *Gli1* is restored with purmorphamine. **(D)** *Vsx2* declines due to *Lhx2* inactivation. *Gli1* is expressed in response to purmorphamine. **(E)** *Gli1* expression declines in the absence of Shh-N (vehicle) whereas *Vsx2* is maintained. *Gli1* is restored with Shh-N. **(F)** *Vsx2* declines due to *Lhx2* inactivation. Similar to purmorphamine, *Gli1* expression is upregulated with Shh-N. **(G)** qPCR-based *Gli1* expression in control and CKO retinal explants at the start of the culture (t=0) and after 24 hr at different concentrations of Shh-N, as determined from a pilot dose response with wild type retinal explants (Supplemental Fig. 6C, Supplemental Table 5). Expression values are relative to the mean control value at t=0 (closed circle). Orange lines extend from t=0 values for the control (upper line) and the CKO (closed triangle, lower line). Note that the value for the 24 hr control in 0 nM Shh-N overlaps with the CKO at t=0. To the right of the graph is the summary table for 2-way ANOVA showing that the main effects (genotype and Shh-N concentration) are significant and interact. See Supplemental Table 5 for statistics including multiple comparisons.

After 24hr in culture, *Gli1* expression was markedly reduced in the untreated explants compared to explants treated with 10 μM purmorphamine (Fig. 6C). Since the RPC gene *Vsx2* was still abundantly expressed (Fig. 6C), the drop in *Gli1* expression in the untreated control explants was not due to RPC loss but instead to a specific reduction in Shh signaling. RGCs exhibited enhanced apoptosis (Supplemental Fig. 6A,B), potentially reducing the availability of endogenous Shh (Wang et al., 2002). We therefore attribute the expression of *Gli1* in the treated control explants to purmorphamine. Interestingly, *Gli1* expression was activated in CKO explants treated with purmorphamine (Fig. 6D). Since the initial level of *Gli1* in the CKO explants was already reduced due to *Lhx2* inactivation, this result indicates that pathway activation at the level of Smo can occur in the absence of Lhx2. We also treated retinal explants with 20 nM Shh-N and obtained similar results (Fig. 6E,F). These observations support the idea that the Shh pathway extending from Smo and downstream signal transduction retains some functionality and that Patched-mediated inhibition of Smo is intact in the CKO retina.

To determine if CKO RPCs respond differently than control RPCs to Shh-N treatment, we assessed *Gli1* expression by qPCR in a pilot dose response experiment with wild type E14.5 retinal explants cultured for 24 hr (Supplemental Fig. 6C). As expected, retinal explants cultured without Shh-N exhibited a strong reduction in *Gli1* expression compared to its level at the start of the experiment (t=0), whereas 3 nM Shh-N was sufficient to maintain signaling, and 10 nM significantly enhanced signaling (Supplemental Fig. 6C; Supplemental Table 5). Based on these observations, we tested the responsiveness of CKO and control explants at 0, 1, and 3 nM Shh-N (Fig. 6G). With all data normalized to *Gli1* expression in control explants at t=0 (solid circle, upper dashed orange line), we made several observations (statistics are listed for all comparisons in Supplemental Table 5). First, when compared to CKO explants at t=0 (closed triangle, lower dashed red line), *Gli1* expression decreased in CKO explants cultured without Shh-N (0nM Shh-N; open triangle,), revealing that endogenous Shh ligand promotes a low level of signaling in the CKO retina. Second, *Gli1* expression dropped in control explants cultured without Shh-N (open circle, 0nM Shh-N) to the same level as the CKO retina at t=0 (closed triangle, lower dashed red line). Third, CKO and control explants exhibited a similar response profile to Shh-N from 0 to 1 nM, but CKO explants lagged behind from 1 to 3 nM. 2-way ANOVA showed significant differences in both genotype and Shh-N concentrations as well as an interaction between both (Fig. 6G; Supplemental Table 5). These data indicate that endogenous Shh is promoting a very low level of signaling in *Lhx2-*deficient RPCs, that *Lhx2-*deficient RPCs can respond to recombinant Shh-N at more physiologically relevant concentrations, but their response is still attenuated compared to *Lhx2*-expressing RPCs.

### Ptch1 inactivation stimulates Shh signaling in the Lhx2-deficient retina but fails to restore retinal development

We next tested if Shh signaling could be stimulated in *Lhx2* CKO RPCs by the simultaneous removal of *Ptch1* function. Conditional *Ptch1* inactivation in the embryonic limb caused increased and ectopic Shh signaling as revealed by elevated readout expression and Shh gain of function phenotypes (Butterfield et al., 2009). Similar to purmorphamine, *Ptch1* inactivation exposes the activity of endogenous Smo in a ligand-independent manner and provides a cell autonomous, *in vivo* test of signaling competence at the level of Smo in *Lhx2*-deficient RPCs. It also allows us to directly test whether Ptch1 is restricting signaling in the *Lhx2* CKO retina and, if so, whether its inactivation improves retinal development.

*Ptch1* CKO, *Lhx2* CKO, and *Ptch1, Lhx2* double CKO (dCKO) retinas were generated with the *Hes1^CreER^* driver (Supplemental Fig. 7A). Retinas were harvested at E15.5 from embryos treated with tamoxifen at E10.75 and E11.5, and recombination of the *Ptch1^flox^* allele was assessed by RT-PCR with primers that amplify both the intact *floxed* and *deleted* transcripts (Supplemental Fig. 7B,C). Non-recombined transcript was detected but its relative abundance was low compared to the deleted transcript. Therefore, all samples were used to measure the relative expression levels of *Gli1, Ptch1, Ptch2,* and *Hhip* by qPCR (Fig. 7A). By 1-way ANOVA, the main effect of genotype on all 4 genes was highly significant (p<0.0001), and for each gene, at least 4 of the 6 pairwise genotype comparisons showed significant differences in expression (Supplemental Table 6). Of those, *Hhip* and *Ptch1* expression were increased in the *Ptch1* CKO compared to control, indicating that *Ptch1* inactivation on its own stimulated Shh signaling. *Gli1* and *Ptch2* expression were decreased in the *Lhx2* CKO, consistent with their high DEG rank in the RNA sequencing data (Fig. 3D). Importantly, the expression levels of all four genes were significantly higher in the dCKO compared to the *Lhx2* CKO, with *Ptch1, Hhip,* and *Ptch2* surpassing control levels (Fig. 7A). These data provide *in vivo* evidence that Lhx2-deficient RPCs retain the competence to signal at the level of Smo, and that Ptch1 is inhibiting signaling in the absence of *Lhx2*.

**Figure 7:**
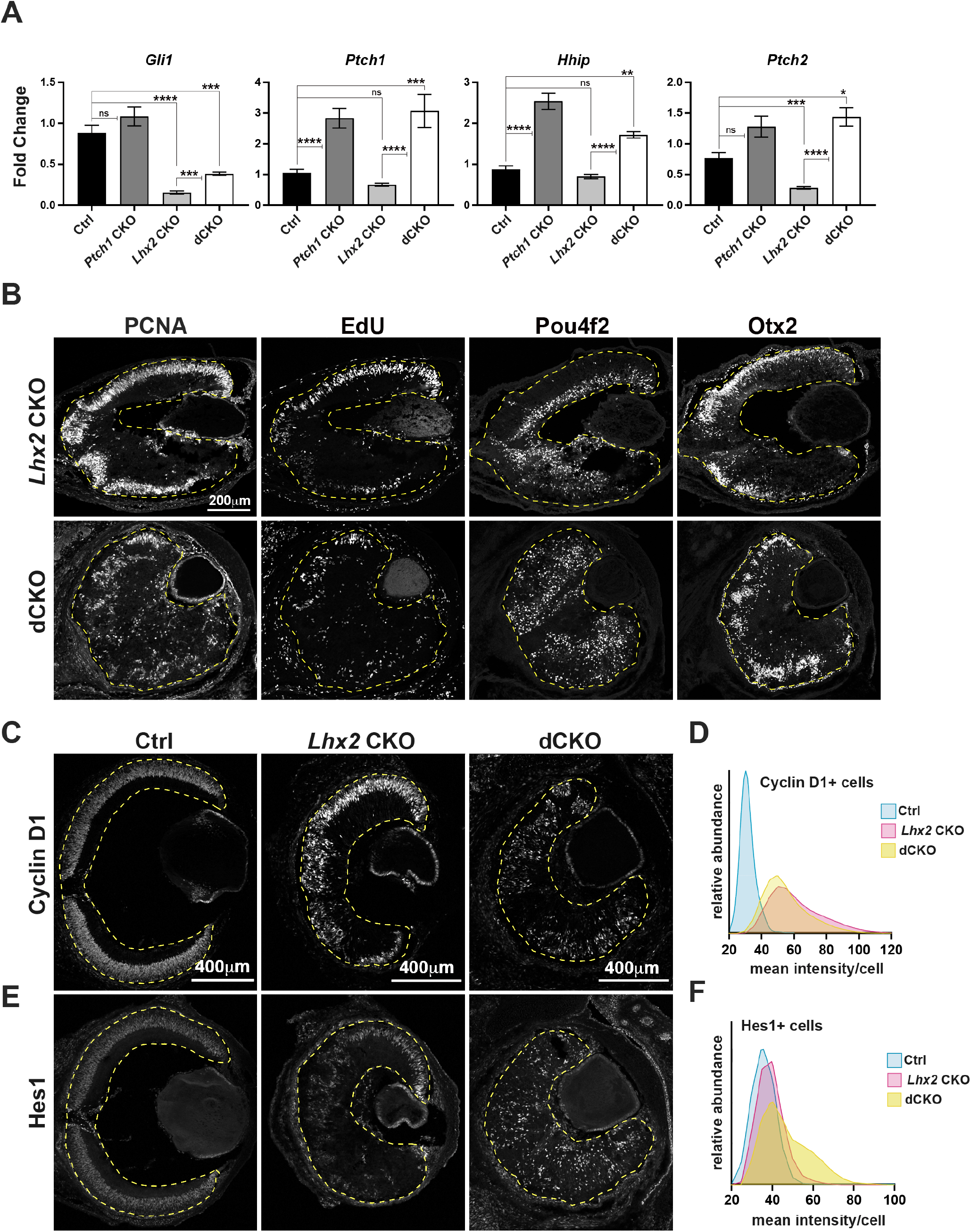
Hh signaling is enhanced in *Lhx2*-deficient RPCs by *Ptch1* inactivation *in vivo*. **(A)** Relative expression of *Gli1*, *Ptch1*, *Hhip*, and *Ptch2* in E15.5 retinas with the following genotypes: control, *Ptch1* CKO, *Lhx2* CKO and *Lhx2; Ptch1* double CKO (dCKO). See Supplemental Fig 7A for breeding scheme and genotypes assigned to control. For each gene, fold change values are relative to a specific control sample set as a reference. Shown are the comparisons for the three mutant genotypes compared to control and for the dCKO compared to the *Lhx2* CKO (ns, not significant; * p_adj_ < 0.05; ** p_adj_ < 0.01; *** p_adj_ < 0.001; **** p_adj_ < 0.0001; See Supplemental Table 6 for statistics and complete multiple comparisons list). **(B)** Expression patterns for PCNA and EdU incorporation to identify RPCs and Pou4f2 and Otx2 to identify nascent RGCs and photoreceptors in E15.5 *Lhx2* CKO (top row) and dCKO (bottom row) retinas. **(C)** Cyclin D1 expression in control, *Lhx2* CKO and dCKO retinas. **(D)** Distribution of Cyclin D1+ cells as a function of the mean fluorescence intensity per cell. Each histogram is normalized to the number of Cyclin D1+ cells within the respective genotype. **(E)** Hes1 expression in control, *Lhx2* CKO and dCKO retinas. **(F)** Distribution of Cyclin D1+ cells as a function of the mean fluorescence intensity per cell. Each histogram is normalized to the number of Cyclin D1+ cells within the respective genotype.

Despite the evidence for increased Shh signaling, the histogenesis defects due to *Lhx2* inactivation did not improve with *Ptch1* inactivation. PCNA staining and EdU incorporation revealed a persistent deficit of RPCs, and the disorganized distribution of Pou4f2+ RGC and Otx2+ photoreceptor precursors were consistent with lamination defects (Fig. 7B). The failure of *Ptch1* inactivation to alleviate, even partially, the *Lhx2* CKO phenotype suggested that Lhx2 also acts downstream of Shh signaling. To assess this further, we examined the expression of Cyclin D1 and Hes1, which are both highly ranked DEGs in the *Lhx2* CKO, are regulated by Shh signaling (Hashimoto et al., 2006; Kenney and Rowitch, 2000; Wall et al., 2009) and are required for retinal neurogenesis (Bosze et al., 2020; Das et al., 2009; Das et al., 2012; Takatsuka et al., 2004). Similar to PCNA and EdU, the abundance of Hes1 and Cyclin D1 expressing cells was similar in dCKO and *Lhx2* CKO retinas (Fig. 7C,E). However, Cyclin D1+ cells appeared brighter in the mutant retinas and was confirmed with fluorescence intensity measurements (Fig. 7D). Hes1+ cells also appeared brighter, but more so in the dCKO retina (Fig. 7F). These changes in cellular fluorescence intensities suggest increased expression, and at least for Hes1, appears to be dependent on *Ptch1* inactivation. These observations suggest that in the absence of Lhx2, elevated Shh signaling due to *Ptch1* inactivation extended to Hes1 but was insufficient to improve retinal development.

### Gas1 and Cdon mediate Lhx2-dependent activation of Shh signaling

While our data show that Lhx2 influences the expression of multiple Shh pathway genes, measurable increases in pathway activity were still achieved in the *Lhx2* CKO with Shh-N or purmorphamine treatment *in vitro*, and *Ptch1* inactivation *in vivo*. These findings indicate that *Lhx2* inactivation did not cause an insurmountable block in the intracellular portion of the pathway and raises the possibility that an additional level of the pathway is dependent on *Lhx2*. Since reduced ligand availability was effectively ruled out, this leaves receptivity to ligand, possibly at the level of co-receptor function. Interestingly, the Shh co-receptors Boc, Cdon, Gas1, and Lrp2 are expressed in the embryonic retina, but Boc and Lrp2 are unlikely candidates because their ranks for differential expression in the RNA-Seq dataset were far below the cutoff (10530 and 11268, respectively) and their genetic inactivation largely spares early retinal neurogenesis (Cases et al., 2015; Fabre et al., 2010). On the other hand, both Gas1 and Cdon qualified as DEGs with DESeq2 ranks of 117 and 1248, respectively, and the ChIP-Seq data supports direct gene regulation by Lhx2 (Fig. 3D,E). *in situ* hybridization and qPCR confirmed their downregulation in the *Lhx2* CKO retina following tamoxifen treatment at E11.5 (Supplemental Fig. 8A, Supplemental Table 3), but their expression at E15.5 in the control retina was limited to the retinal periphery. This is not unexpected since Gas1 and Cdon are negatively regulated by Shh signaling in other tissues (Allen et al., 2007; Tenzen et al., 2006), but their restricted expression at E15.5 made it difficult to determine the extent of their dependence on Lhx2 in the retina. We therefore examined their expression patterns from E11.5 – E13.5, the interval encompassing pathway activation (Fig. 8A, Supplemental Fig. 8B). *Gas1* was more restricted to the peripheral retina but *Cdon* was broadly expressed at E11.5, resolving to the peripheral retina by E13.5. *Cdon* expression decreased in a complementary manner to *Gli1* upregulation as revealed by *Gli1* mRNA and by β-Galactosidase (β-Gal) expression from the *Gli1^lacz^* allele. However, *Cdon* downregulation appeared to be ahead of the central to peripheral wave of *Gli1* expression and had a closer complementarity with *Atoh7*, a neurogenic bHLH gene transiently expressed in RPCs that functions as an RGC competence factor (Brown et al., 2001; Brzezinski et al., 2012; Prasov and Glaser, 2012) (Supplemental Fig. 8B). Whether this reflects a novel mode of *Cdon* regulation is unclear, but the complementarity with *Cdon* and *Gli1* expression is consistent with downregulation upon pathway activation.

**Figure 8:**
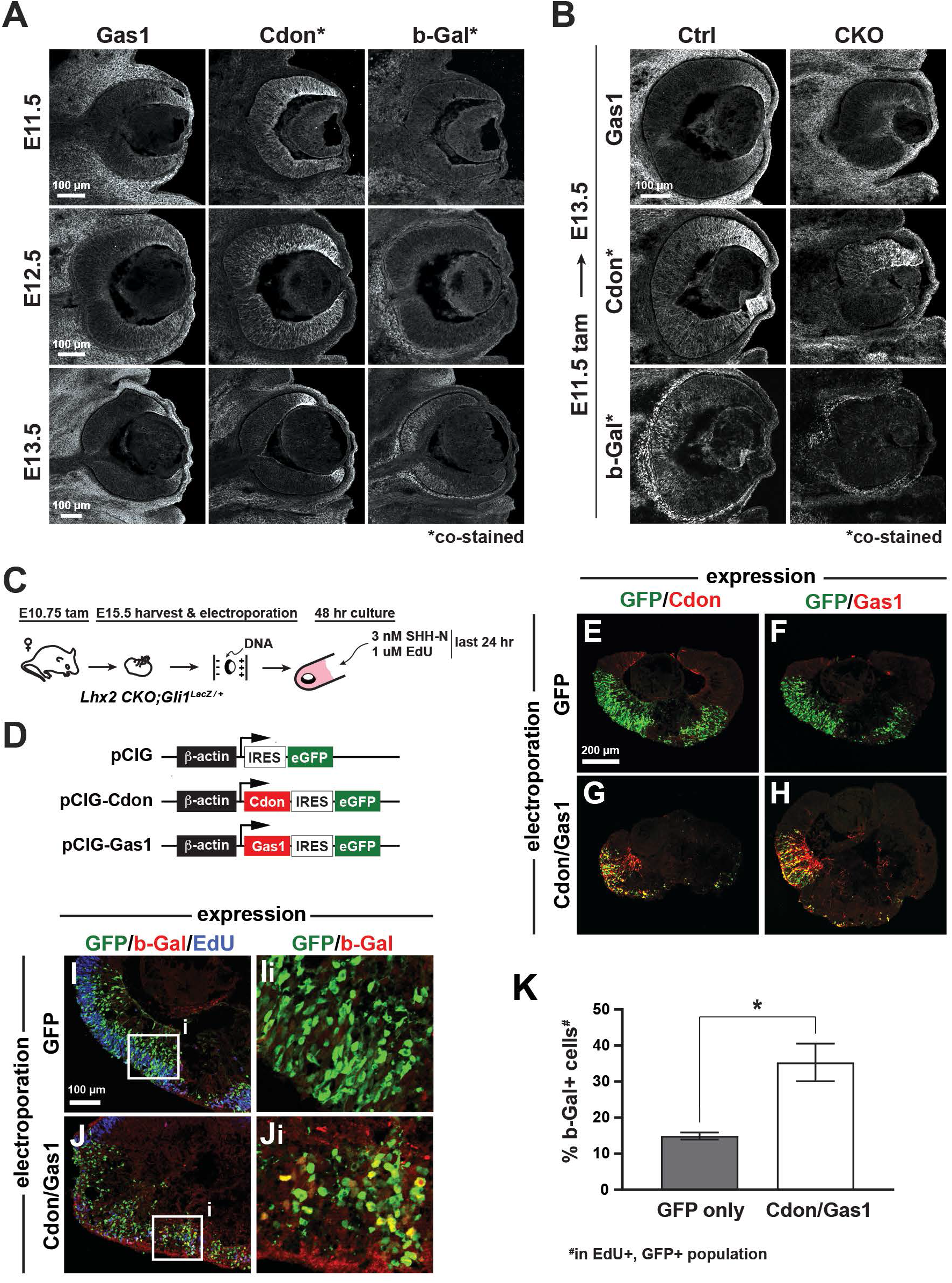
Cdon and Gas1 are functional downstream targets of Lhx2. **(A)** Temporal expression patterns of Gas1, Cdon, and β-Gal at E11.5, E12.5, and E13.5 in *Gli1^lacz/+^* mice. Cdon and β-Gal were detected on the same tissue sections, Gas1 on adjacent sections (also in B). **(B)** Expression of Gas1, Cdon, and β-Gal in Ctrl and *Lhx2* CKO; *Gli1^lacz/+^* eyes at E13.5 following tamoxifen treatment at E11.5. **(C)** Experimental design for *ex vivo* electroporation and explant culture. *Lhx2* CKO; *Gli1^LacZ/+^* explants were electroporated at the beginning of the culture. 3nM Shh-N and 1uM EdU were added after 24hr and cultured for an additional 24hr. **(D)** DNA constructs used for electroporation. pCIG served as the control and *pCIG-Cdon* and *pCIG-Gas1* were co-electroporated. **(E, F)** Explants were electroporated with *pCIG* and co-stained for GFP and Cdon (E) or Gas1 (F). **(G, H)** Explants were co-electroporated with *pCIG-Cdon* and *pCIG-Gas1* and co-stained for GFP and Cdon (G) or Gas1 (H). **(I, J)**. Electroporated explants were co-stained for GFP, β-Gal, and EdU. Insets (Ii, Ji) show GFP and β-Gal staining only. **(K)** Quantification of the percentage β-Gal+ cells in the EdU+, GFP+ cell populations from GFP (control) and Cdon/Gas1 electroporated (* p<0.05; See Supplemental Table 7 for statistics).

We next examined *Gas1* and *Cdon* expression in the CKO at E12.5 (Supplemental Fig. 8C) and E13.5 (Fig. 8B) following tamoxifen treatment at E10.5 and E11.5, respectively. At E12.5, the Lhx2 target *Vsx2* was downregulated as were *Gas1* and *Cdon*, consistent with Lhx2 promoting their expression, but it was too early to assess effects on *Gli1* (Supplemental Fig. 8C). At E13.5, Gas1 and Cdon proteins were downregulated and β-Gal was not detected in the CKO (Fig. 8B). These observations support the idea that Lhx2 promotes the expression of *Cdon* and *Gas1* to confer signaling competence to RPCs. Interestingly, *Cdon* expression persisted to some extent in the dorsal CKO retina at both ages similar to what we observed for *Gli1* (Fig. 8B, Supplemental Fig. 8C, Supplemental Fig. 1B). Whether this reflects mechanistic differences in Shh signaling or how Lhx2 regulates Shh signaling across the retina is unclear.

To determine if *Gas1* and *Cdon* were sufficient to restore signaling, we returned to the *ex vivo* culture paradigm (Fig. 8C). *Lhx2* CKO; *Gli1^LacZ^* explants were electroporated with *pCIG* or *pCIG-Gas1* and *pCIG-Cdon* at the start of the culture (Fig. 8D). After 24 hr, 3nM Shh-N was added for an additional 24 hr. Since electroporation is not cell type specific, 1μM EdU was also added to label proliferating RPCs (Fig. 8C). Gas1 (red) and Cdon (red) expression were not detected in CKO explants transfected with *pCIG* alone (GFP, green) (Fig. 8E,F), but were readily detected in explants co-transfected with *pCIG-Gas1* and *pCIG-Cdon* (Fig. 8G,H). Triple labeling for GFP (green), EdU (blue), and β-Gal (red) revealed a significant increase in *Gli1* reporter positive cells in explants co-transfected with *pCIG-Gas1* and *pCIG-Cdon* compared to explants transfected with *pCIG* only (Fig. 8I-K; Supplemental Table 7). These data support the hypothesis that *Lhx2* confers signaling competence in RPCs through promoting *Cdon* and/or *Gas1* expression.

### Lhx2 promotes Shh signaling after Gas1 and Cdon downregulation

Although re-expressing *Cdon* and *Gas1* increased *Gli1* expression in the absence of *Lhx2*, it remains that reaching or maintaining the appropriate level of signaling could still depend on regulation of other Shh pathway genes by Lhx2. Since *Cdon* and *Gas1* are downregulated by E13.5, inactivating *Lhx2* after E13.5 provided an opportunity to test this. Tamoxifen was administered at E14.5 and retinas collected at E17.5 for qPCR to assess changes in relative gene expression of *Gli1, Gli2*, *Ptch2,* and *Smo* (Fig. 9A; Supplemental Table 3). *Lhx2* downregulation was highly efficient and accompanied by the predicted drop in *Vsx2*. Interestingly, *Gli1, Gli2*, *Ptch2,* and *Smo* were also reduced in the CKO retina. These changes were not due to developmental disruptions caused by *Lhx2* inactivation because early retinal neurogenesis is largely spared with tamoxifen treatment by E13.5 (Gordon et al., 2013). Furthermore, the reductions in *Gli2, Ptch2,* and *Smo* are not strictly due to reduced *Gli1* activity because their expression levels were not significantly altered in the *Gli1* KO retina at E15.5 (Fig. 9B; Supplemental Table 3). From this, we conclude that Lhx2 promotes Shh signaling at more than one point in the pathway with a measurable influence on signaling that is independent of Cdon and Gas1.

**Figure 9:**
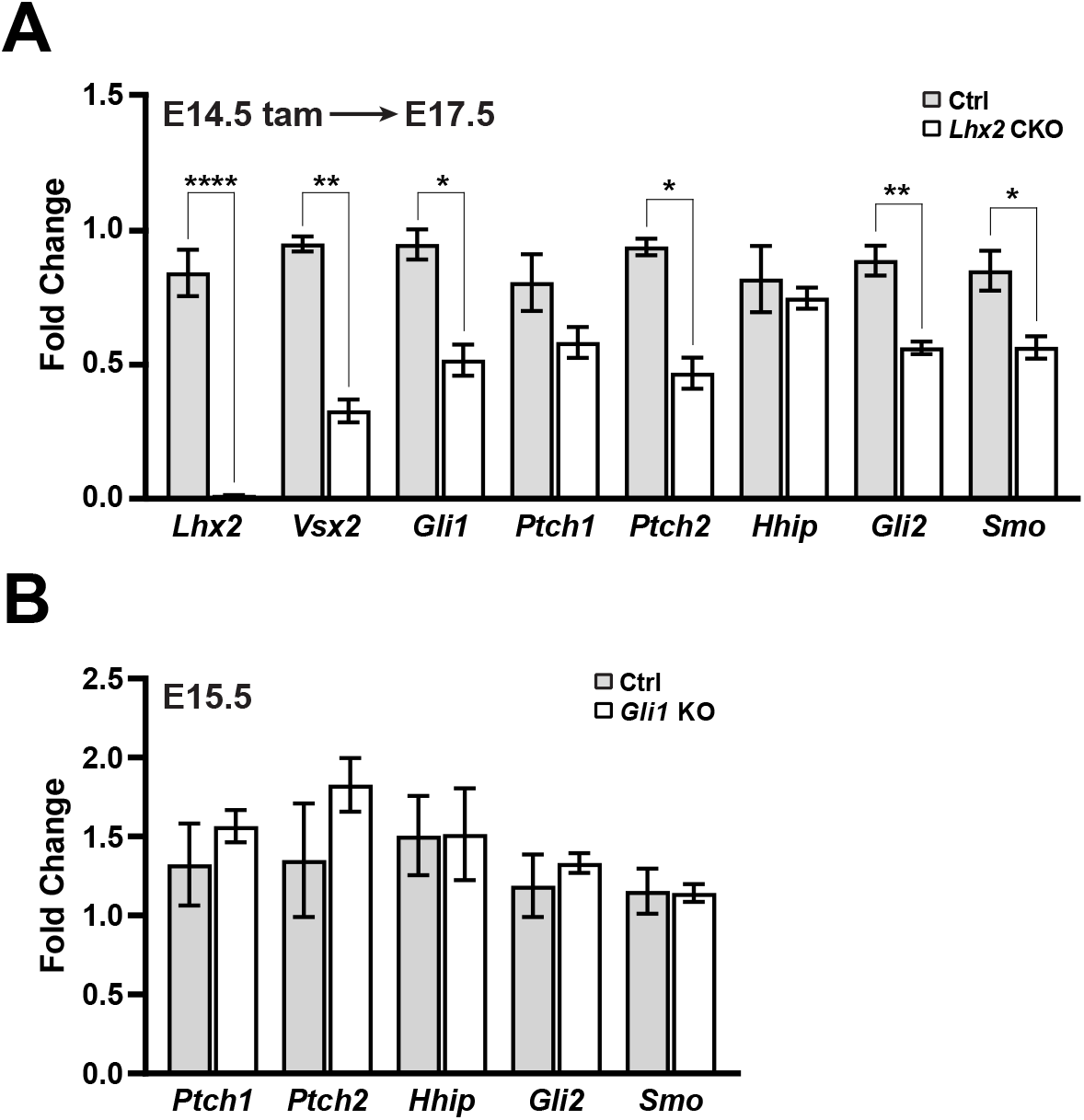
Lhx2 is required for sustained Hh signaling. **(A)** qPCR-based expression for *Lhx2, Vsx2, Gli1, Ptch1, Ptch2, Hhip, Gli2,* and *Smo* from E17.5 control and *Lhx2* CKO retinas following tamixofen treatment at E14.5. For each gene, fold change values are relative to a specific control sample set as a reference. Only significant comparisons are noted (* p_adj_ < 0.05; ** p_adj_ < 0.01; **** p_adj_ < 0.0001; see Supplemental Table 3 for statistics) **(B)** qPCR-based expression for *Ptch1, Ptch2, Hhip, Gli2,* and *Smo* from E15.5 control (*Gli1^LacZ/+^*) and *Gli1* KO retinas. For each gene, fold change values are relative to a specific control sample set as a reference. None of the comparisons were significant (See Supplemental Table 3 for statistics).

## Discussion

### Lhx2 is a multilevel modulator of the Shh pathway

Lhx2 and Shh signaling are essential regulators of vertebrate retinal development that function at multiple stages. Here, we present evidence supporting a model in which Lhx2 promotes the expression of multiple genes in the Shh pathway that allows for the timely activation and sustained levels of signaling during early retinal neurogenesis by conferring signaling competence to RPCs (Fig. 10A). Directly supporting Lhx2’s role in this regard, we show that endogenous Shh ligand was functional and available in the *Lhx2*-deficient retina. Rather, Lhx2 supports ligand reception by what is likely direct regulation of expression of the co-receptors *Cdon* and *Gas1* and efficient signaling by promoting *Smo* and *Gli2* expression, possibly through indirect mechanisms (Fig. 10B). As revealed by Shh-N treatment or *Ptch1* inactivation in the *Lhx2-*deficient retina, *Gli1* expression remains Shh-dependent, but in the specific context of the *Lhx2*-deficient retina, the reduced expression of *Gli1* could combine with reductions in *Gli2* and *Smo* to negatively impact sustained Shh signaling. Since Lhx2 regulates a broad repertoire of RPC genes at the epigenetic and transcriptional level (Gueta et al., 2016; Yun et al., 2009; Zibetti et al., 2019), Lhx2 could also link Shh signaling to its downstream transcriptional targets as suggested by the failure of *Ptch1* inactivation to alleviate, even partially, the phenotypic consequences of *Lhx2* inactivation on early retinal neurogenesis. Based on the sum of our observations, we propose that Lhx2, in addition to promoting Shh signaling in RPCs, integrates the pathway into the program of early retinal neurogenesis.

**Figure 10:**
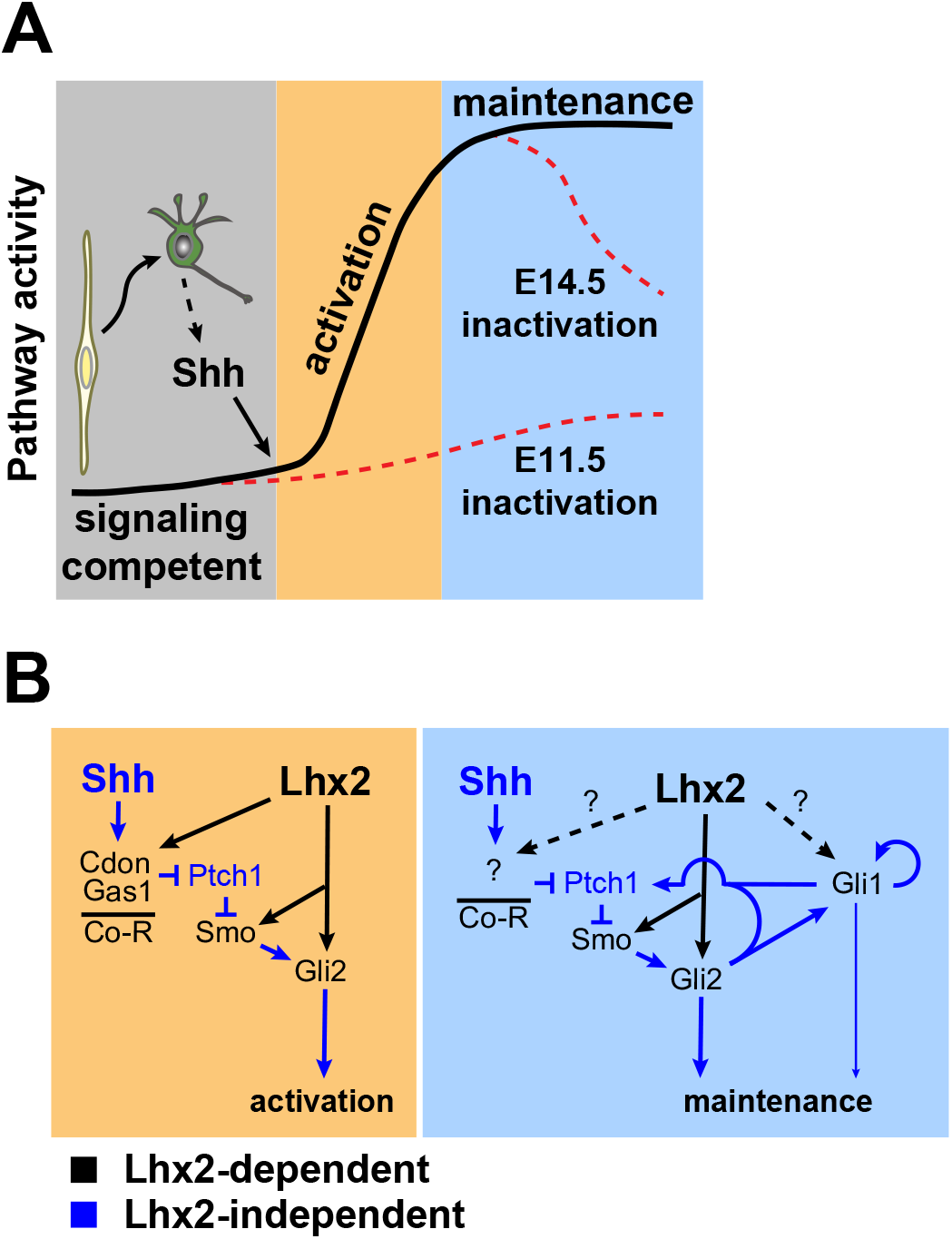
Models of Lhx2’s interactions with Hh signaling during embryonic retinal neurogenesis. **(A)** At the cellular level, Lhx2 promotes signaling competence in RPCs and once neurogenesis begins, RGCs produce ligand leading to pathway activation. This is revealed by Lhx2 inactivation at E11.5. Lhx2 also promotes the correct level of signaling in RPCs as evidenced by the drop in pathway readout gene expression when Lhx2 is inactivated at E14.5. **(B)** Mechanistically, Lhx2 promotes signaling competence and efficient activation by promoting the expression of the coreceptors (Co-R) for ligand reception, Smo for signal transduction, and Gli2 for target gene activation. During the maintenance phase, Lhx2 promotes signaling again by promoting Smo and Gli2. Lhx2 may also regulate other coreceptors, Gli1, or other factors to promote efficient signaling.

### Cdon and Gas1 are central mediators of Lhx2’s role in promoting Shh pathway activation

Our data indicate that Lhx2 interacts with the Shh pathway in a complex manner, but the links to Cdon and Gas1 are likely to transmit the largest impact on pathway activation. These linkages are revealed by the rapid loss of Cdon and Gas1 expression after *Lhx2* inactivation and by ChIP-seq data showing Lhx2 binding at the *Cdon* and *Gas1* loci. Upstream of Cdon and Gas1, we show that ligand availability was intact, and immediately downstream, purmorphamine treatment and *Ptch1* inactivation stimulated signaling in the *Lhx2*-deficient retina, as did Cdon and Gas1 overexpression. These observations all point to a role for Lhx2 in ligand reception upstream of Smo and Ptch1, a function fulfilled by Shh co-receptors such as Cdon and Gas1.

This seemingly straightforward requirement for Cdon and Gas1 in Lhx2-mediated Shh signaling contrasts with other identified roles for Shh co-receptors in early eye and retinal development. Prior to the onset of retinal neurogenesis, Cdon is expressed in the optic vesicle and functions in a manner consistent with both promoting and inhibiting Shh signaling (Gallardo and Bovolenta, 2018). Cdon’s positive role is revealed by Cdon KO mice, which exhibit a range of phenotypes consistent with absent or impaired Shh signaling, including holoprosencephaly (HPE) and Septo Optic Dysplasia (SOD) (Bae et al., 2011; Cavodeassi et al., 2018; Kahn et al., 2017; Zhao et al., 2012). While these early and severe phenotypes would normally preclude an assessment of its later role in RPCs, retinal development occurs in *Cdon* mutant mice that evade the HPE and SOD phenotypes (Zhang et al., 2009). In this context, the *Cdon* mutant phenotypes align well with retinal-specific Shh loss of function mutants, and similar to what we show here for *Lhx2* inactivation, *Cdon*-deficient RPCs fail to express *Gli1* even though *Shh* is still expressed (Kahn et al., 2017). Cdon’s inhibitory effect on Shh signaling was revealed in chick and zebrafish optic vesicles, where it limits the range of Shh signaling through ligand sequestration in signaling-*incompetent* optic neuroepithelial cells (Cardozo et al., 2014). An inhibitory role for another co-receptor, Lrp2, occurs in the nascent ciliary epithelium, where it is proposed to mediate endocytic clearance of Shh and prevent binding to Ptch1(Christ et al., 2015). The nascent ciliary epithelium arises from the peripheral edge of the retinal neuroepithelium, the same location where Gas1 and Cdon were downregulated at E15.5 following *Lhx2* inactivation at E11.5 (Supplemental Fig. 8A). Since Shh signaling is ectopically activated in this domain in *Lrp2* mutant mice (Christ et al., 2015), a primary role for Lrp2 in the developing ciliary epithelium could be to prevent Cdon and Gas1 from activating the pathway. Based on the sum of these findings, we propose that Lhx2 sets up the signaling competence of RPCs by promoting Cdon expression throughout the retina and Gas1 expression in the peripheral retina. This is countered by Lrp2 in the far retinal periphery, where suppression of Shh signaling is required for ciliary epithelium development.

How Gas1 relates to Shh signaling in early eye development is less clear. During optic vesicle patterning, *Gas1* is expressed in the nascent retinal pigment epithelium (RPE) and in *Gas1* KO mice, the predominant phenotype is an RPE to neural retina transformation in the ventral optic cup (Lee et al., 2001). This does not phenocopy the effects of directly disrupting Shh signaling, where inhibition reduces the proximal and ventral domains of the optic vesicle, and overactivation expands them (Amato et al., 2004; Cavodeassi et al., 2018; Kim and Lemke, 2006). The most similar outcomes were observed in temporally controlled cyclopamine treatments in *Xenopus* embryos, where ventral RPE differentiation was disrupted, but a retinal fate transformation was not reported (Perron et al., 2003). It therefore remains unclear if the requirement for Gas1 during early eye and retinal development is related to Shh signaling beyond what we propose here. Direct analysis of Gas1’s requirement in RPCs could help to clarify this.

### Lhx2 deficiency reveals potential differences in RPC receptivity to endogenous and recombinant Shh-N

Given the importance of the co-receptors in pathway activation, the upregulation of *Gli1* in the *Lhx2* CKO explants treated with recombinant Shh-N was unexpected (Fig. 6). One possibility is residual co-receptor activity after *Lhx2* inactivation. Indeed, *Cdon* and *Gli1* expression persisted in the dorsal retina for a few days after tamoxifen treatment, albeit in disrupted (*Cdon*) and diminished (*Gli1*) patterns. However, 20 nM Shh-N treatment induced *Gli1* across the CKO retina (Fig. 6F) and although we did not track the axial orientation of the explants, this continuous pattern was consistently observed, making it unlikely that we were only sampling dorsal retina.

Another possibility is that recombinant Shh-N bypassed the requirement for co-receptors in pathway activation. Supporting this, Ptch1 and Shh-N can form ternary complexes (Ptch1:Shh-N:Ptch1) that are capable of promoting signaling although with lower efficiency compared to complexes containing coreceptors (i.e. Cdon:Shh-N:Ptch1) (Beachy et al., 2010; Qi et al., 2019; Qi and Li, 2020; Qi et al., 2018). Another consideration is that endogenous Shh is post-translationally lipidated with cholesterol and palmitoyl moieties, key adducts for efficient signaling (Manikowski et al., 2018). Following secretion by ligand-producing cells, endogenous Shh is kept soluble while in transit to responding cells by forming a ‘lipid-shielding’ complex with SCUBE proteins (Tukachinsky et al., 2012). The Shh:Scube2 complex initially forms a ternary complex with Cdon or Boc on the responding cell. The Shh:Scube2 complex is then transferred to Gas1, which releases Shh from Scube2 and allows Ptch1 to bind Shh to initiate signaling (Wierbowski et al., 2020). In contrast, the recombinant Shh-N used here is not lipid modified and is soluble in its native form. It can directly bind Ptch1, obviating the need for Gas1 to disengage Shh-N from a complex with Scube. In the context of the *Lhx2* CKO retina, then, endogenous Shh could have been rendered less efficient than recombinant Shh-N because of the differential requirement for the co-receptors to receive endogenous Shh and remove Scube.

### Lhx2 regulates multiple pathway components to achieve the optimal level of signaling

Since *Cdon* and *Gas1* are downregulated in the retina by E14.5 but Shh signaling persists, the importance of Lhx2’s control over their expression is only relevant for pathway activation. However, *Lhx2* inactivation at E14.5 also reduced pathway activity. It’s possible that Lhx2 regulates another co-receptor such as Lrp2 to sustain Shh signaling (Fig. 10B) but this is unlikely since the *Lrp2* CKO retina does not have a phenotype at this stage (Cases et al., 2015). Rather, we propose that Lhx2’s role in sustained signaling is co-receptor independent and through regulation of other cell intrinsic pathway components. Our data does not support the existence of a single, essential pathway component that is under strong Lhx2 regulation, but instead suggests that Lhx2 exerts a more subtle regulation of multiple components of the Shh pathway (Fig. 10B), and we identified *Smo*, *Gli2*, and *Gli1* as candidate Lhx2-dependent genes. Genetic reductions in *Smo* or *Gli2* on their own do not exhibit haploinsufficiency (Mo et al., 1997; Sakagami et al., 2009), but their combined reduced expression could result in pathway sensitization that if strong enough, could cause a synthetic haploinsufficiency. Supporting this, *Gli2* heterozygous mice are phenotypically normal, but exhibit greater teratogenic sensitivity to vesmodegib, a small molecule Smo inhibitor, as compared to their wild type littermates (Heyne et al., 2016). If the reductions in *Smo* and *Gli2* expression were still not sufficient to reduce signaling, additional changes in other pathway components such as *Gli1* could have shifted the balance. Although we did not uncover evidence of Lhx2 binding in or near the *Gli1* locus, it is possible that Lhx2 exerts some control over *Gli1* expression (Fig. 10B). Importantly, an interaction of this nature would have to be context specific since *Gli1* inactivation, on its own, has minimal effect on development and expression of Shh target genes (Fig. 9B) (Bai et al., 2002; Furimsky and Wallace, 2006; McNeill et al., 2012; Park et al., 2000; Wall et al., 2009). Thus, subtle or partial regulation of multiple pathway genes by Lhx2 could confer optimal levels of signaling, first during activation in conjunction with strong regulatory input to the coreceptors, and during sustained signaling, after co-receptor downregulation. This could be especially important in the retina where *Shh* expression is comparatively lower than in other tissues but exhibits a qualitatively similar level of *Gli1* expression (Sigulinsky et al., 2021).

In sum, we propose that Lhx2 regulates the expression of multiple pathway components required for optimal Shh signaling in RPCs, both during and after pathway activation. Although Lhx2 does not regulate ligand availability at a functional level, it does limit the production of RGCs (Gordon et al., 2013). Because RGCs are the primary source of retinal Shh ligand, Lhx2 acts in an indirect but semi-autonomous manner to limit ligand expression in the retina. Furthermore, since Lhx2 is an essential RPC transcription factor, it also likely acts to link Shh signaling to downstream targets as suggested by the persistent phenotypic severity of the *Ptch1*, *Lhx2* dCKO retina. Through this molecular and cellular circuitry, Shh signaling is tailored by Lhx2 to meet the demands of early retina formation.

## Materials and Methods

### Animals

All procedures and experiments involving animals were approved by the Institutional Animal Care and Use Committees at the University of Utah and Vanderbilt University, and set forth in the Association for Research in Vision and Ophthalmology (ARVO) Statement for the Use of Animals.

Single night matings were set up in the late afternoon and females were checked for plugs the next morning. Embryonic age determinations were based on plug date with noon designated as E0.5, by weight determinations, and by morphological criteria (Theiler, 2013). Pregnant dams were euthanized with a Euthanex EP-1305 CO_2_ delivery system according to manufacturer’s instructions and AALAC guidelines. Uteri were removed and embryos retrieved in Hanks Buffered Saline Solution (HBSS) supplemented with 20 mM HEPES and 6 mg/ml glucose at room temperature. The embryos were removed from the placenta and extraembryonic membranes, rapidly euthanized by decapitation with surgical scissors. Embryonic tissue was collected for PCR genotyping and whole heads or dissected eyes were processed as needed for each analysis (see below).

### Genetics and breedings

All alleles used in this study were generated previously and are listed in Supplemental Table 8. *Hes1^CreERT2^; Lhx2^flox/+/-^; Rosa26^ai14^* strains are described in Gordon et al (2013). The combinatorial strains needed for experiments utilizing the *Ptch1^flox^* and *Gli1^lacz^* alleles were produced through successive rounds of strategic breeding beginning with crosses to *Hes1^CreERT2^; Lhx2^+/-^* and/or *Lhx2^flox/flox^; Rosa26^ai14/ai14^* mice. To generate embryos for analysis, all breedings were done in which the male carried the *Hes1^CreERT2^* allele in a heterozygous state. Embryos that underwent Cre recombination were selected at time of dissection by detection of tdTomato or eGFP using an epi-fluorescence stereomicroscope. When possible, CKO and control embryos were segregated by the presence or absence of microphthalmia, respectively, and subsequently verified by PCR genotyping.

All genotypes were verified by PCR based genomic DNA genotyping. Primers are listed in Supplemental Table 9. Deletion of *Ptch1* exon 3 was confirmed by RT-PCR using the same retinal RNA preparations used for qPCR. All PCR conditions are available on request.

### Tamoxifen treatments

Tamoxifen (Sigma T5648) was dissolved in corn oil (Sigma C8267) and administered by oral gavage to pregnant dams at dosages ranging between 0.15 – 0.2 mg per gram body weight. For *Ptch1* CKO and dCKO matings, two doses of 0.1mg per gram body weight were administered 24 hr apart. Treatment times are noted in the text for each experiment.

### Western Blots

Western blots were done as described in Ringuette et.al. (2016) with the following modifications. For E15.5 samples, 10 retinas (5 embryos) per genotype were pooled and 6 retinas (3 pups) were pooled for P0 samples. 50 ug protein was loaded per lane. Primary antibodies used were mouse anti-Gli1 (1:3000, Cell Signaling Technologies) and mouse anti-γ-Tubulin (1:1000, Sigma, Cat#T6557), the latter serving as the loading control. Detection was done with donkey anti-mouse IgG horseradish peroxidase at 1:5000 (1:5000, Millipore Cat#AP308P) and Luminata Crescendo Western HRP substrate (Millipore Cat#WBLUR0100). Blots were probed for Gli1 first, stripped, and reprobed for γ-Tubulin.

### Immunohistology and *in situ* hybridization

Embryo heads, eyes, or explants were fixed in 4% PFA/1xPBS at 4°C from 45 minutes (explants) to 2hr (heads), rinsed in PBS, cryoprotected in 20% sucrose/PBS, embedded in OCT (Sakura Finetek, Torrance, CA), and stored at -80°C. Frozen tissues were sectioned on a Leica CM1950 cryostat at a thickness of 12 μm.

Primary antibodies are listed in Supplemental Table 10. Primary antibodies were followed with species-specific secondary antibodies conjugated to Alexa Fluor 488, 568, or 647 (Invitrogen/Molecular Probes, Eugene, OR). EdU was detected using Click-iT EdU Cell Proliferation kit for imaging (Thermo Fisher Scientific). Nuclei were stained with 4,6-diamidino-2-phenylindole (DAPI; Fluka). Panels showing fluorescence-based protein detection are single scan confocal images obtained with a Fluoview 1000 confocal microscope (Olympus) or Zeiss LSM710 confocal microscope equipped with 20X objective.

*In situ* hybridization was performed as previously described (Gordon et al., 2013; Sigulinsky et al., 2008). Probes used in this study were digoxigenin-labeled anti-sense probes against *Atoh7, Cdon, Gas1, Gli1, Shh,* and *Vsx2*.

### RNA sequencing

Total RNA was isolated from flash frozen retinal tissue using QIAshredder columns and RNeasy Micro kit (Qiagen, Cat#79654 and 74004). Libraries were constructed with Illumina TruSeq RNA Library Kit V2 with poly(A) selection, and 50 cycle single end sequencing was done on the Illumina Hi-Seq 2000 platform. RNA data alignment was performed by TopHat 2 (Trapnell et al., 2009) on MM10 reference genome followed by gene quantification into FPKM using Cufflinks (Trapnell et al., 2010). Additional read count per gene was generated using HTSeq (Anders et al., 2015). FASTQ files are deposited at GEO repository number GSE172457.

### Bioinformatics

#### RNA-seq data analysis

Differential gene expression analysis was performed with DESeq2 (version 1.21.4) (Love et al., 2014). Features (i.e. genes) were removed that contained 0 counts across all samples or when fewer than 3 samples had normalized counts greater than or equal to 20. The Canonical Pathways algorithm in the Ingenuity Pathway Analysis Suite (Qiagen) was performed on the DEGs bounded by an upper FDR cutoff of 0.001 from the DESeq2 analysis.

#### ChIP-seq and ATAC-seq data analysis

FASTQ files from E14 Lhx2 ChIP-seq, wildtype ATAC-seq and Lhx2 CKO ATAC-seq were obtained from GEO repository number GSE99818 (Zibetti et al., 2019). Sequencing adaptors were trimmed away using NGmerge (Gaspar, 2018). Bowtie2 was used for read alignment on the GRCm38/mm10 mouse genome (Langmead and Salzberg, 2012). deepTools2 was used to generate bigwig files of the sequencing data which were then visualized on the UCSC genome browser (Ramírez et al., 2016) and https://genome.ucsc.edu/index.html). Peak calling was performed using Genrich (available at https://github.com/jsh58/Genrich). High mapping-quality reads were kept (MAPQ > 10) and mitochondrial aligned reads and PCR duplicate read were filtered out. The blacklisted regions in mouse were excluded from peak regions (https://github.com/Boyle-Lab/Blacklist/blob/master/lists/mm10-blacklist.v2.bed.gz) (Amemiya et al., 2019). ChIPseeker was used to annotate Lhx2 ChIP-seq peaks and identify the closest gene to the peak or genes within 10kb of the peak (Yu et al., 2015). The gene lists were then used for KEGG pathway enrichment using the enrichKEGG function in clusterProfiler (Yu et al., 2012). For ATAC-seq peak calling, the -j option was used to set up Generich in ATAC mode. A consensus list of peaks was generated by merging all the ATAC peaks sets and filtering peaks that were reproducible in at least 2 samples. DESeq2 was used to identify differential accessible regions (DARs) between wildtype and Lhx2 CKO ATAC datasets from the consensus peak list (Love et al., 2014).

### Quantitative reverse transcription PCR (qPCR)

Relative changes in gene expression were determined with the delta-delta-Ct method (DDCt). *Gapdh* served as the internal control gene for the initial normalization (DCt values). For E14.5 *Lhx2* CKO and control retinas, DDCt values were generated by normalizing DCt values to the mean DCt value of the control samples for each gene. In the Shh-N dose response experiments, DDCt values were generated by normalizing DCt values to the mean DCt value of the control samples at the start of the experiment (t=0). For samples from the E15.5 combinatorial *Lhx2* and *Ptch1* CKO and control retinas, E17.5 *Lhx2* CKO and control retinas, and E15.5 *Gli1* KO and control retinas, DDCt values were generated by normalization to a single control sample that was designated as a *reference*. A sample reference was used to control for potential batch-effect variations due to reactions being run on different plates and/or days. Data is presented in graphs as the *fold change* in gene expression based on RQ values (2^-DDCt^).

#### Sybr Green assays

Sybr Green-based qPCR was done on samples from E14.5 *Lhx2* CKO and control retinas, and for the dose response experiments with recombinant Shh-N in retinal explants. Total RNA was isolated with RNeasy Micro and cDNA was synthesized with the SuperScript III First Strand Synthesis System (Invitrogen, Cat#18080-051). qPCR was done on a BioRad CFX96 Real Time PCR system with SsoAdvanced Universal SYBR Green Supermix (BioRad, Cat#170-8841) and primers detecting *Vsx2, Ascl1, Sox2, Lhx2, Pax6, Lin28b, Prtg, Gas1, Cdon, Boc, Gli, Shh, Smo*, and *Gapdh* (Supplemental Table 9). All primer pairs were first validated for reaction efficiency and specificity.

#### TaqMan assays

TaqMan-based qPCR was done on samples from E15.5 combinatorial *Lhx2* and *Ptch1* CKO and control retinas, E17.5 *Lhx2* CKO and control retinas, and E15.5 *Gli1* KO and control retinas. Total RNA was isolated using TRIzol (Thermo Fisher Scientific, Cat#15596026) and cDNAs were synthesized using SuperScript IV VILO master mix (Thermo Fisher Scientific, Cat# 11766051). qPCR was done on QuantStudio 3 Real Time PCR Systems (Thermo Fisher Scientific) with the TaqMan gene expression Master Mix (Thermo Fisher Scientific, Cat# 444557) and TaqMan gene probes for *Gli1*, *Hhip*, *Ptch1*, *Ptch2*, *Gapdh* (Supplemental Table 9).

#### Data analysis

Data collection was done with BioRad CFX manager and ABI QuantStudio 3 package for Sybr Green- and TaqMan-based qPCR, respectively. Graphing and hypothesis testing were done with Microsoft Excel (version 16.43) and GraphPad Prism (version 9.0). Descriptive statistics for RQ values (sample number (n), mean, standard deviation (SD), standard error from mean (SEM)), hypothesis tests, testing parameters, and test results are provided in Supplemental Tables 3, 5 and 6. All hypothesis testing was done on DDCt values.

### Cocultures to test endogenous ligand availability

#### Preparation and passaging of the NIH3T3 open-loop responder cell line

Cells were maintained up to 14 passages in 60mm^2^ petri dishes and cultured in 1x DMEM supplemented with 10% calf serum. 8.8 × 10^5^ cells were plated in 60mm^2^ dish for passaging (3-4 days). For experiments, 1.6 × 10^5^ cells were seeded into each well of the 24 well plates and grown to >90% confluence before beginning the experiment (2-3 days).

#### Dose-response testing of open loop cells

Shh-N was added to the culture medium at the doses indicated in the text, which were empirically determined in pilot experiments. A 50% medium exchange with Shh-N at the original concentration was done at 2 DIV.

#### Co-cultures

Whole retina was dissected away from other ocular tissues and placed onto the underside of a 0.4μm Biopore insert (Millipore, PICM3050) with the apical layer of the retina in contact with the Biopore membrane (unless noted otherwise). The insert was turned over and placed into a well of a 24-well plate containing a confluent monolayer of NIH3T3 open-loop responder cells. Contact between the basal surface of the retina and the cell monolayer was achieved by removing the filter insert support legs and applying gentle manual pressure to the insert for approximately 4 seconds. The culture medium was raised to the bottom of the insert (approximately 300 ul) with 60 -100 ul media changes per day.

#### Live imaging, post-image processing, and data analysis

Wide-field epifluorescence images were taken with a Nikon DS Qi2 camera on a Nikon TE-200 inverted microscope with a 10x objective for the dose response cultures and a 4x objective for the cocultures. All comparable images were captured with similar camera settings and illumination. Potential daily variations in illumination and sample background fluorescence were managed with post capture background subtraction (see next paragraph).

Image processing and quantification were done with ImageJ (version 2.1.0). In brief, images were processed to obtain regions of interest (ROI) masks for identification of mCitrine+ objects (nuclei). All images were first processed with background subtraction (rolling ball algorithm) and bit depth conversion from 14 to 8 prior to mask generation and measurement. Masks were then generated through thresholding, binarization, and watershedding. Masks were then applied to their respective images for object counts and pixel intensity calculations (average mean intensity per object, integrated densities). Objects smaller than 90 µm^2^, an empirically estimated measure of a nucleus, were excluded. Objects that passed this filter were retained for quantification and considered to represent cells on the basis of 1 nucleus/cell and designated as such. Objects with larger areas were attributed to overlapping mCitrine+ nuclei. These larger objects were also retained for analysis since their exclusion would exacerbate underrepresentation of cell counts and fluorescence intensities in the conditions with the highest number and brightest objects. Some inflation of fluorescence intensity was likely due to overlapping nuclei although the use of average mean intensities per object as a measure should have substantially reduced the impact of this confounding variable.

Graphing and hypothesis testing were done with Microsoft Excel (version 16.43) and GraphPad Prism (version 9.0). Descriptive statistics (sample number (n), mean, standard deviation (SD), standard error from mean (SEM)), hypothesis tests, testing parameters, and test results are provided in Supplemental Table 4.

### Organotypic suspension cultures

Whole retina and lens were dissected away from other ocular tissues (explants) and placed into a 14ml round bottomed snap cap tube with 1ml of retina culture medium (RCM) and rotated at 15 RPM on a carousel with an axis of rotation at 30° above horizontal. Explants were incubated at 37°C in a humidified, 5% CO_2_ atmosphere. RCM is composed of 1x DMEM/F12 (US Biological, Cat# D9807-05), 1% fetal bovine serum (Thermo Fisher Scientific, Cat#16140071), 6 mg/ml glucose (Sigma, Cat# G7528), 0.1% NaHCO_3_ (Thermo Fisher Scientific, Cat#25080-094), 50 mM HEPES (Thermo Fisher Scientific, Cat# 15630-080, 1 mM glutamax (Thermo Fisher Scientific, Cat# 35050-061), 1x N2-plus supplement (R&D Systems, Cat#AR003), and 1x penicillin/streptomycin (Thermo Fisher Scientific, Cat# 15070-063).

At the end of the culture period, explants were rinsed in PBS. For *in situ* hybridization or immunohistology, explants were fixed in 4%PFA and prepared for cryostorage (see above). For qPCR, lens tissue was removed, and retinas were snap frozen in liquid N_2_, and stored at -80 °C until use.

#### Purmorphamine and Shh-N treatments

Purmorphamine (EMD Chemicals, Cat# 540220) was dissolved in 100% DMSO at 2.5mM and stored at -20°C. Explants were treated with 10µM purmorphamine or vehicle (0.4% DMSO). E. coli-derived human N-terminal modified (C24II) fragment of Sonic Hedgehog (Shh-N; R&D Systems, Cat#1845-SH/CF) was reconstituted in PBS at 10µM and stored at -80°C. Explants were treated with Shh-N at the concentrations indicated. When possible, multiple breeding pairs were set per single night mating to increase sample size. Explants were cultured for 24hr.

See qPCR methods for data collection and analysis.

#### Ex vivo electroporation

*Lhx2* CKO, *Gli1*^lacz/+^ embryos were identified by phenotyping and rapid PCR genotyping. Explants were transferred into sterile PBS without Ca^2+^ and Mg^2+^. 1.5 µl of DNA (3 µg/µl) in 30% glycerol with methyl green was pipetted onto the apical retinal surface, and electroporated with a BTX ECM830 (5 × 50ms pulses at 50V, 250ms intervals). One explant per animal received the GFP control plasmid (*pCIG*) and the other received an equimolar 50:50 mixture of *pCIG-Cdon* and *pCIG-Gas1*. Explants were transferred to 1 ml of RCM and cultured. 3nM Shh-N and 1 µM EdU were added after 24hr, and the cultures were maintained for an additional 24 hours. Explants were fixed for 1 hr in 4% PFA and cryopreserved until use.

Sectioned explants were stained for GFP to identify transfected cells, EdU to identify proliferating cells (RPCs), and β-Gal to identify *Gli1* expressing cells. Cells were first scored for the co-localization of GFP and EdU. Once completed, GFP, EdU double positive cells were scored for β-Gal expression. Counting was done on at least three sections per explant, and when possible, at least 100 GFP/EdU double positive cells were counted per section. Graphing and hypothesis testing were done with Microsoft Excel (version 16.43) and GraphPad Prism (version 9.0). Descriptive statistics (counts per explant, sample number (n), mean, standard deviation (SD), standard error from mean (SEM)), hypothesis tests, testing parameters, and test results are provided in Supplemental Table 7.

## Acknowledgements

We thank Michael Lewis (Baylor College of Medicine, Houston TX) for the *Ptch1^flox^; Rosa26^mT/mG^* mice, Guoqiang Gu (Vanderbilt University) for the *pCIG-GFP* plasmid, Ben Allen for the *pCIG-Cdon* and *pCIG-Gas1* plasmids, Joe Brzezinski for training on the electroporation technique, JP Cartailler (Creative Data Solutions) for bioinformatics. We also thank members of the Levine laboratory at the University of Utah (Anna Clark, Crystal Sigulinsky) and Vanderbilt University Medical Center (Allison Klinger, Dianna Rowe, Mahesh Rao, Amanda Leung,) for their technical and intellectual support. We also thank Sabine Fuhrmann and the Fuhrmann laboratory for sharing reagents, technical expertise, and intellectual input.

## Author Contributions

**EML:** Conceived the study, performed experiments, data analysis, wrote and edited ms

**XL:** performed experiments, data analysis, edited ms

**PG:** Conceived the study, performed experiments, data analysis, wrote and edited ms

**JG:** performed experiments, data analysis, edited ms

**AF:** performed experiments, data analysis, edited ms

**CS:** data analysis, edited ms

**SB:** data analysis, edited ms

**RR:** performed experiments

**VW:** data analysis, edited ms

**PL:** data analysis, edited ms

## Supplemental Figure Legends

**Supplemental Figure 1: (A, B)** *in situ* hybridizations for *Gli1* expression in E16.5 Ctrl (**A**) and CKO (**B**) eyes following tamoxifen treatment at E12.5. **(C, D)** *in situ* hybridizations for *Shh* expression in E16.5 Ctrl (**C**) and CKO (**D**) eyes following tamoxifen treatment at E12.5. Dashed boxes reveal locations of close-up images (i, ii).

Abbreviations: D, dorsal; V, ventral; DCL, differentiated cell layer; NBL, neuroblast layer

**Supplemental Figure 2: ATAC-Seq and Lhx2 ChIP-Seq tracks for selected Hh pathway genes within the FDR cutoff for differential expression (0.001).** ATAC-Seq and Lhx2 ChIP-Seq tracks from E14.5 RPCs. Differentially accessible chromatin regions (DARs) in *Lhx2* CKO RPCs compared to control are indicated by red (open) and blue bars (closed). Lhx2 ChIPSeq peak calls are indicated by grey bars.

**Supplemental Figure 3: ATAC-Seq and Lhx2 ChIP-Seq tracks for selected Hh pathway genes outside the FDR cutoff for differential expression (0.001).** ATAC-Seq and Lhx2 ChIP-Seq tracks from E14.5 RPCs. Differentially accessible chromatin regions (DARs) in *Lhx2* CKO RPCs compared to control are indicated by red (open) and blue bars (closed). Lhx2 ChIPSeq peak calls are indicated by grey bars.

**Supplemental Figure 4: Responder cells express mCitrine when placed into close proximity with the basal surface of the retina.**

**(A)** Schematic cross-section of embryonic retina shows that the apical surface is comprised mainly of RPCs and developing photoreceptors (cones until ∼E15.5, and a mix of rods and cones thereafter) and the basal surface is comprised of RGCs, the source of retinal SHH. Astrocytes and endothelial also reside on the basal surface (not shown) **(C)** E18.5 wild type retinas were cultured in opposite orientations such that responder cells came into contact with the basal or apical surfaces of the retina. mCitrine expression was robustly induced in responder cells in contact or close proximity to the basal surface.

**Supplemental Figure 5: Single channel images**

Single channel fluorescence and bright field images for the cocultures shown in Fig. 5B.

**Supplemental Figure 6: Markers of recombination, apoptosis, and RGCs after 24hr, and dose response to estimate physiological range for recombinant Shh-N(C24II) protein.**

**(A)** Control and **(B)** CKO explants treated with vehicle or 10 nM purmorphamine for 24 hr. Left column: tdTomato expression from heterozygous *ai14* allele reveals CreER activity throughout the explants. Middle column: Caspase 3 immunofluorescence (green) reveals extent of apoptotic cells. Nuclei are stained with DAPI (blue). Right column: Pou4f2 immunofluorescence (green) marks RGCs. **(C)** Relative *Gli1* expression in wild type E15.5 retinas treated with increasing concentrations of recombinant Shh-N. Data points are relative to the mean expression value from retinas harvested at the start of the experiment (t=0; orange line). Means and standard deviation are shown, and results are from a pilot experiment. See Supplemental Table 5 for statistics.

**Supplemental Figure 7: Genetics of *Ptch1* and *Lhx2* inactivation and validation of *Ptch1* recombination.**

**(A)** Overview of genetics and tamoxifen treatment. **(B)** Schematic of *Ptch1* locus encompassing the regions containing exons 2-6. Exon 2 forward (ex2F) and Exon 6 reverse (ex6R) were used for RT-PCR to detect unrecombined and recombined transcripts from the flox and CKO alleles, respectively. Agarose gel shows most samples were recombined (CKO) although some unrecombined (flox) was detected. All samples were used for qPCR.

**Supplemental Figure 8: *in situ* hybridizations.**

**(A)** Expression of *Gas1, Cdon,* and *Gli1* at E15.5 after tamoxifen treatment at E11.5. Boxes in *Gas1* and *Cdon* panels mark locations of insets, which correspond to the retinal periphery, where *Gas1* and *Cdon* are expressed. Arrowheads in Ctrl panels show the restricted peripheral expression that is absent in the CKO. Bracket in *Gli1* CKO panel reveals low, but persistent, expression in the dorsal retina. **(B)** Temporal expression patterns of *Gas1*, *Cdon*, *Gli1*, and *Atoh7* from E11.5 – E13.5. Dashed lines in E12.5 and E13.5 images show that temporal downregulation of *Cdon* mRNA has a tighter spatial complementarity with *Atoh7* mRNA accumulation than *Gli1.* **(C)** Expression of *Vsx2*, *Gli1*, *Gas1*, and *Cdon* at E12.5 after tamoxifen treatment at E10.5. The presence of *Gli1* in the periocular mesenchyme indicates that our staining was sufficient to detect expression. Circles in *Gas1* images show regions of *Gas1* expression in peripheral retina that is lost upon *Lhx2* inactivation. Bracket in *Cdon* CKO image shows absence of *Cdon* in ventral domain, and arrowheads point to persistent *Cdon* expression. Abbreviations: c, cornea; cb, ciliary body; le, lens; onh, optic nerve head; nr, neural retina; pom, periocular mesenchyme.

## Supplemental Tables

**Supplemental Table 1: Differential gene expression analysis of RNA-seq data with DESeq2.**

**Supplemental Table 2: KEGG pathways identified by Ingenuity Pathway Analysis and clusterProfiler.**

**Supplemental Table 3: Statistics and tests for multiple pairwise comparisons of qPCR-based measurements of gene expression (2 genotypes).**

**Supplemental Table 4: Statistics and test results for experiments with open-loop responder cells.**

**Supplemental Table 5: Statistics and test results for Shh-N dose responses in retinal explants.**

**Supplemental Table 6: Statistics and tests for multiple pairwise comparisons of qPCR-based measurements of gene expression (4 genotypes).**

**Supplemental Table 7: Statistics and test results for cell counts following electroporation in *Lhx2* CKO, *Gli1^Lacz/+^* retinal explants.**

**Supplemental Table 8: Alleles**

**Supplemental Table 9: Primers**

**Supplemental Table 10: Antibodies**

