## Supplemental Figures 1-8 for "Lhx2 is a progenitor-intrinsic modulator of Sonic Hedgehog signaling during early retinal neurogenesis"

### Supplemental Figure 1

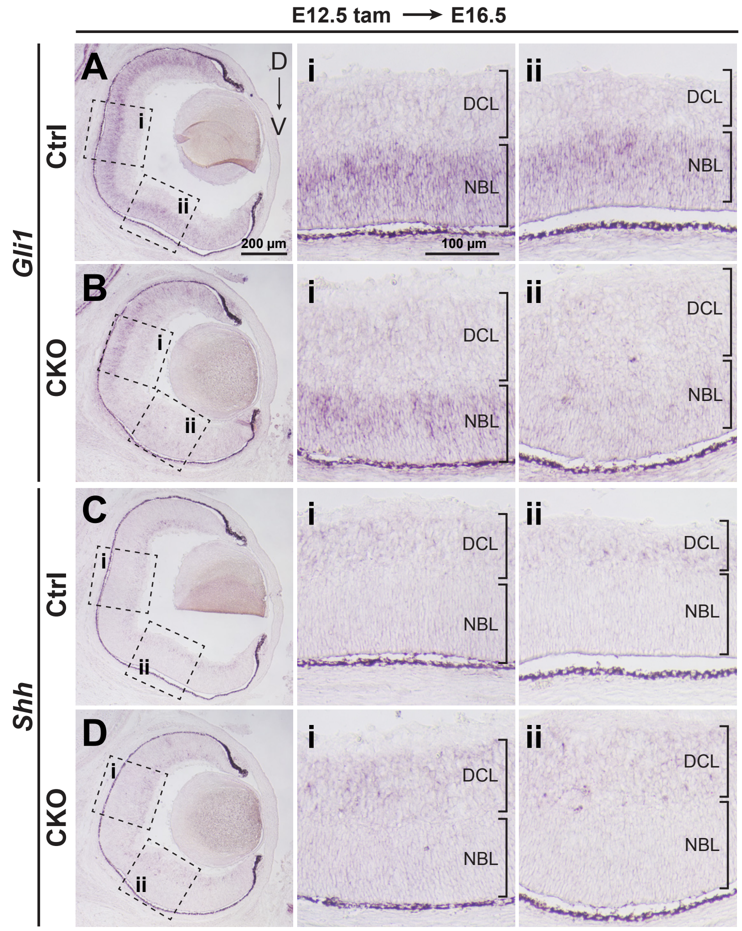

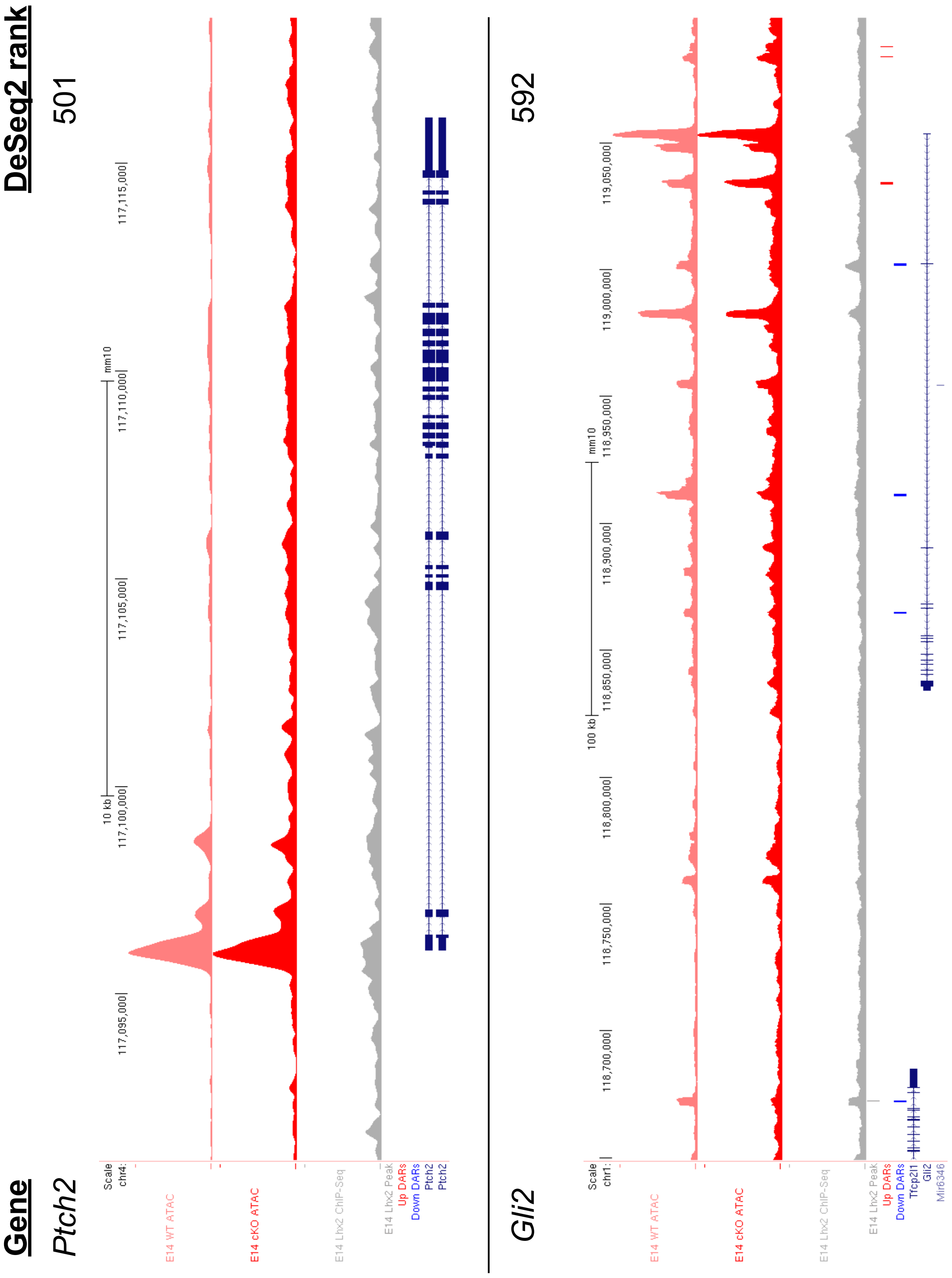
Gene
DeSeq2 rank
Ptch2
501
Scale
chr4:
10 kb
mm10
117,095,000
117,100,000
117,105,000
117,110,000
117,115,000
E14 WT ATAC
E14 cKO ATAC
E14 Lhx2 ChIP-Seq
E14 Lhx2 Peak
Up DARs
Down DARs
Ptch2
Ptch2
Gli2
592
Scale
chr1:
100 kb
mm10
118,700,000
118,750,000
118,800,000
118,850,000
118,900,000
118,950,000
119,000,000
119,050,000
E14 WT ATAC
E14 cKO ATAC
E14 Lhx2 ChIP-Seq
E14 Lhx2 Peak
Up DARs
Down DARs
Tfcp2l1
Gli2
Mir6346

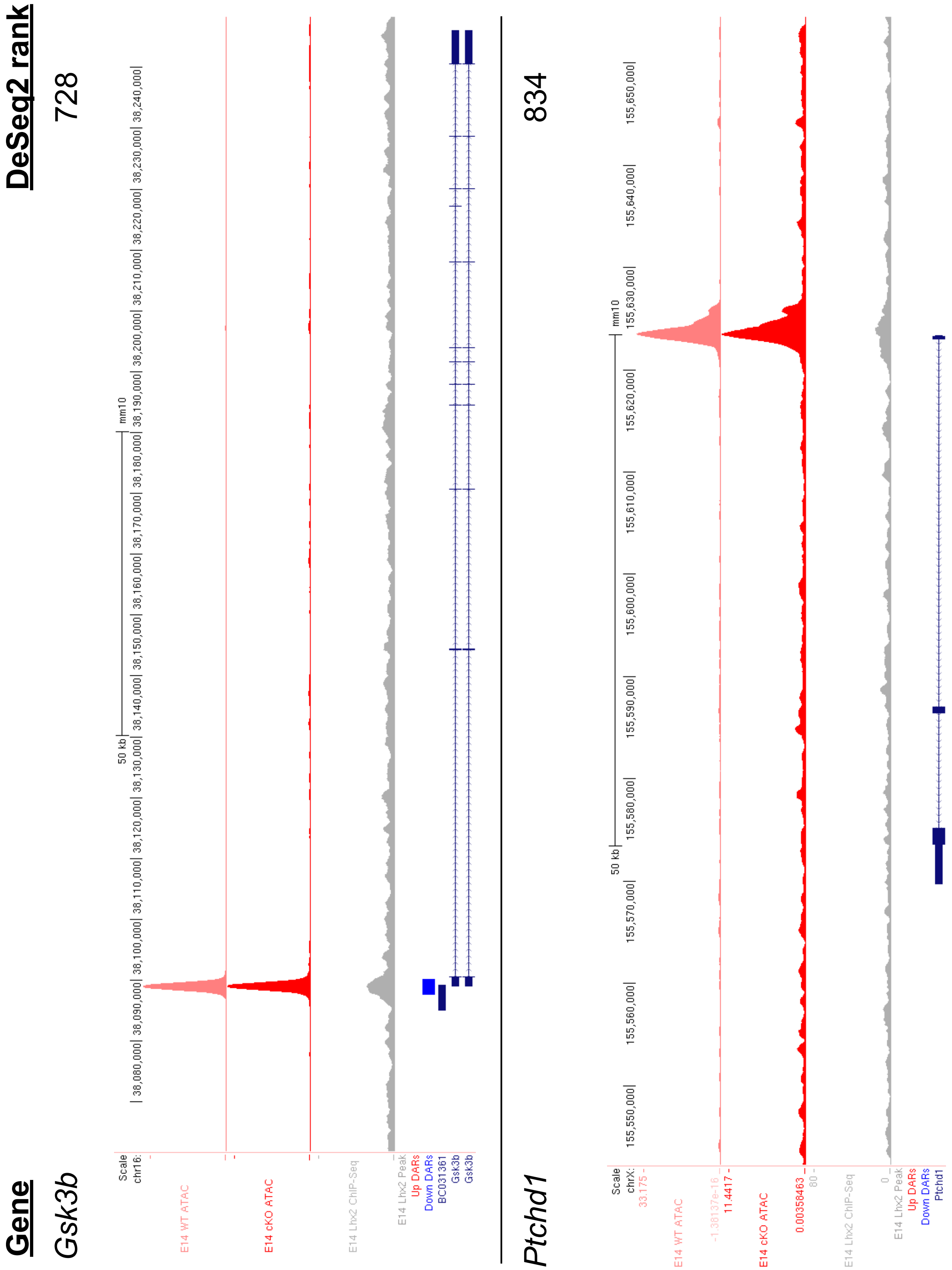

Gene
DeSeq2 rank
Gsk3b
728
Scale
chr16:
50 kb
mm10
E14 WT ATAC
E14 cKO ATAC
E14 Lhx2 ChIP-Seq
E14 Lhx2 Peak
Up DARs
Down DARs
BC031361
Gsk3b
Gsk3b
Ptchd1
834
Scale
chrX:
33.175 -
-1.38137e-16 _
11.4417 -
0.00358463 _
80 -
0 _
E14 WT ATAC
E14 cKO ATAC
E14 Lhx2 ChIP-Seq
E14 Lhx2 Peak
Up DARs
Down DARs
Ptchd1

Supplemental Figure 2 (p.3 of 4)

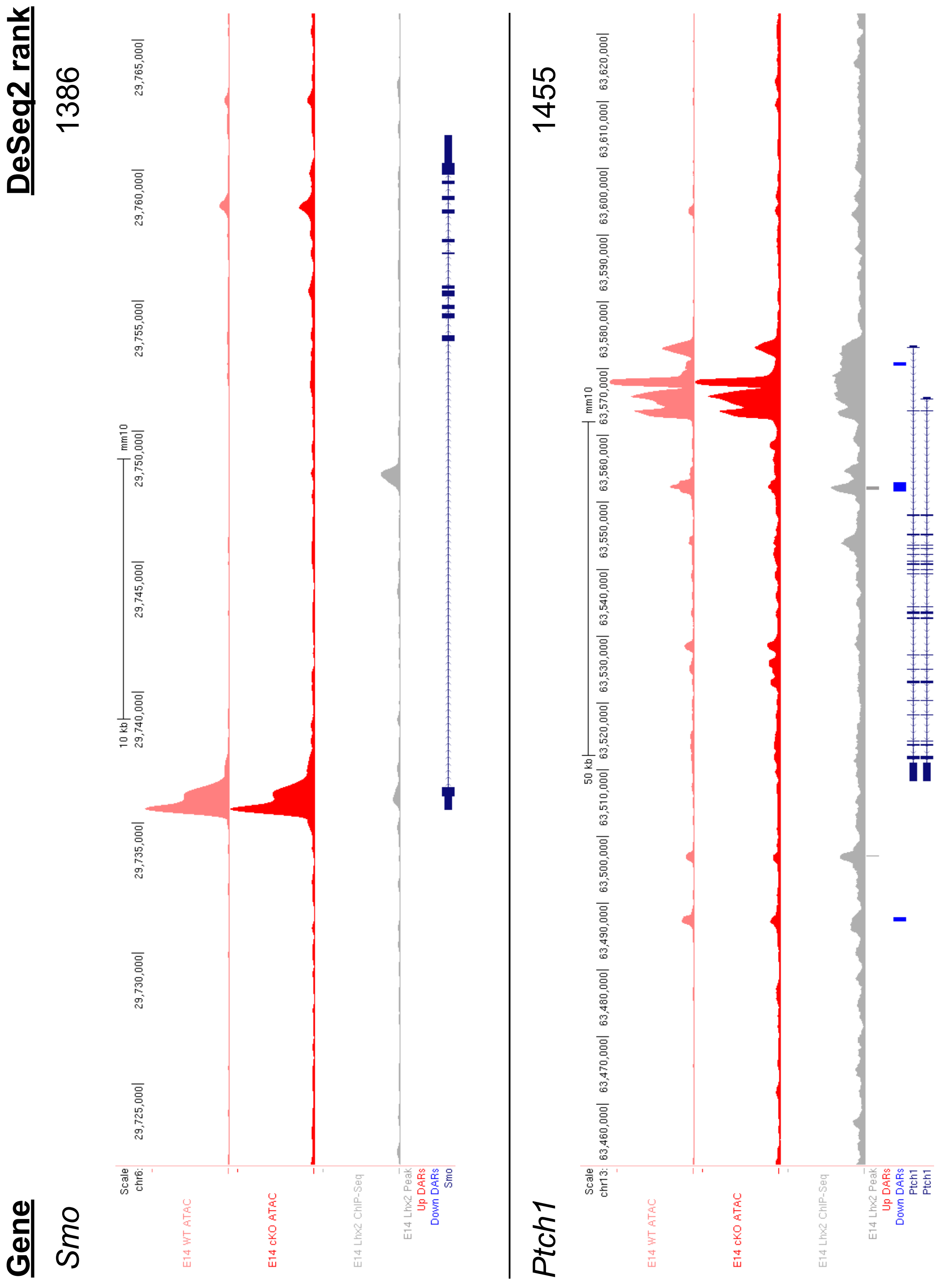

Supplemental Figure 2 (p.4 of 4)

**Gene** | **DeSeq2 rank**
---|---
*Gli3* | 1532

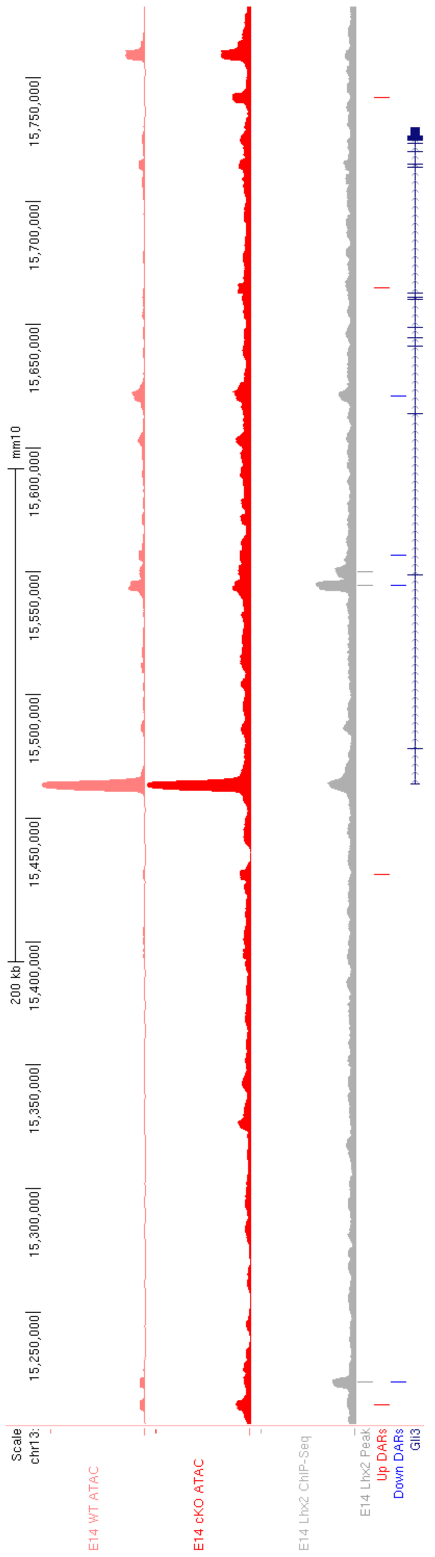

Supplemental Figure 3 (p. 1 of 1)

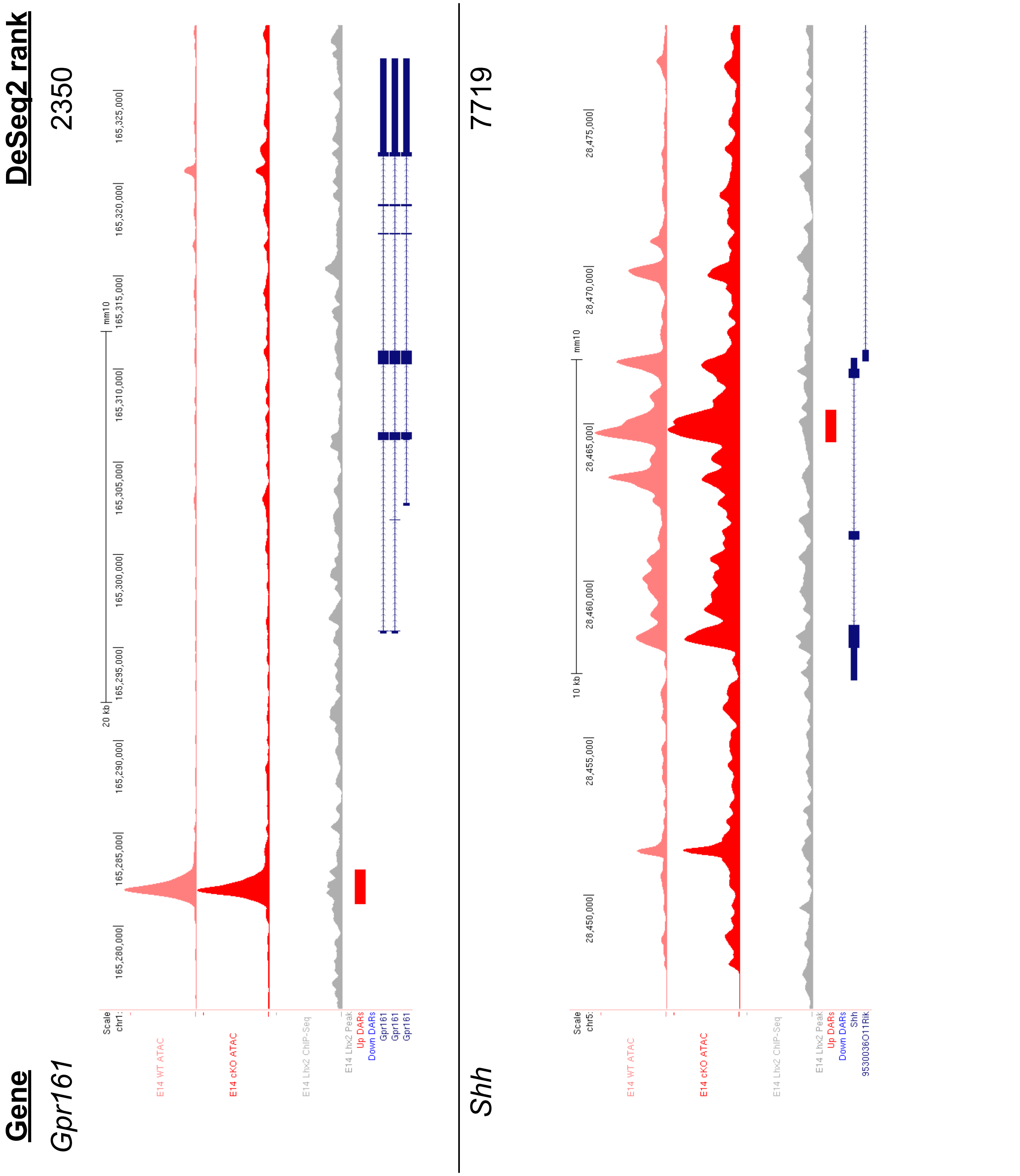

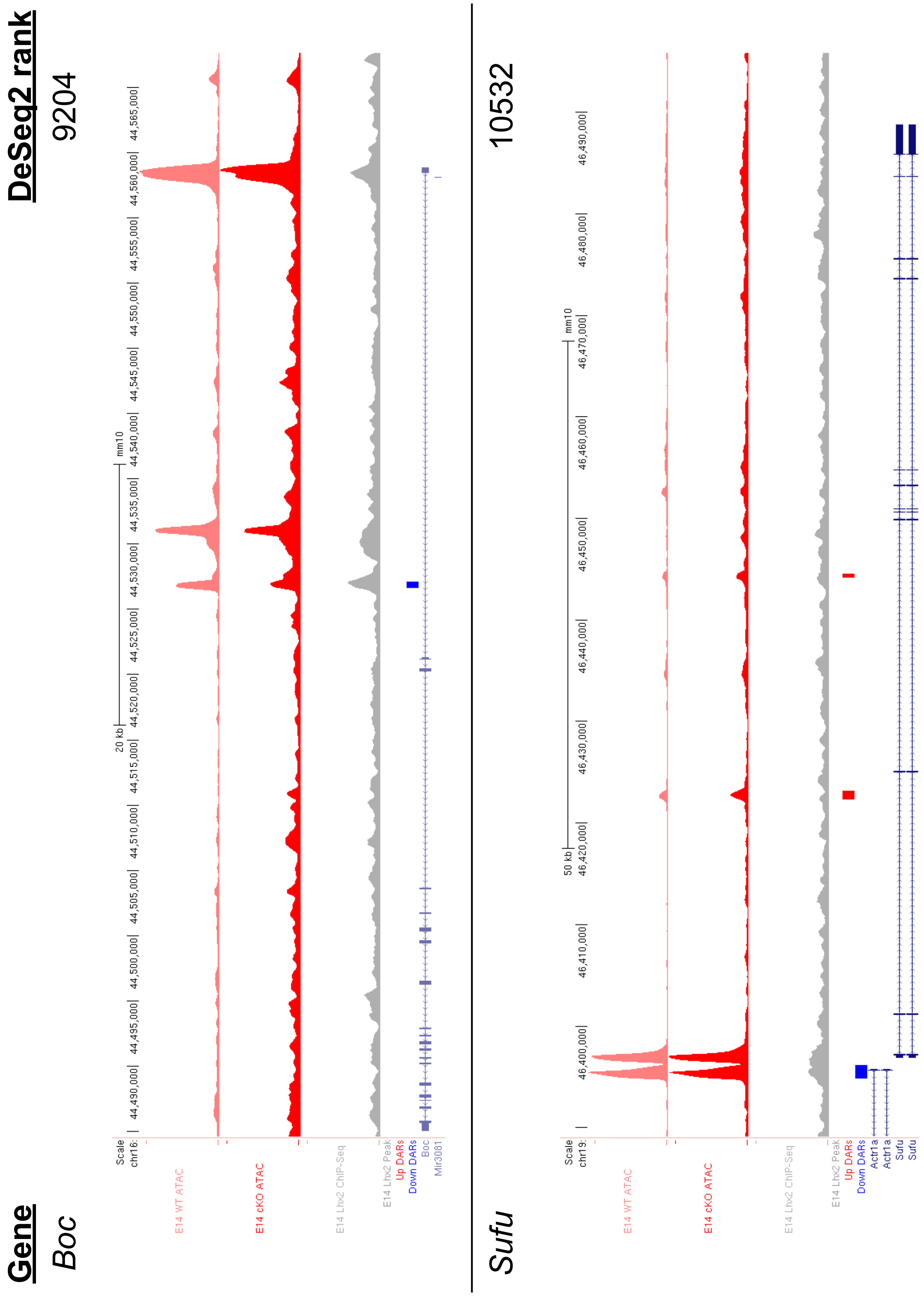
Gene
DeSeq2 rank
Boc
9204
Sufu
10532
Scale
chr16:
20 kb
mm10
44,490,000|
44,495,000|
44,500,000|
44,505,000|
44,510,000|
44,515,000|
44,520,000|
44,525,000|
44,530,000|
44,535,000|
44,540,000|
44,545,000|
44,550,000|
44,555,000|
44,560,000|
44,565,000|
E14 WT ATAC
E14 cKO ATAC
E14 Lhx2 ChIP-Seq
E14 Lhx2 Peak
Up DARs
Down DARs
Boc
Mir3081
chr19:
50 kb
46,400,000|
46,410,000|
46,420,000|
46,430,000|
46,440,000|
46,450,000|
46,460,000|
46,470,000|
46,480,000|
46,490,000|
Actr1a
Actr1a
Sufu
Sufu

Supplemental Figure 3 (p. 3 of 3)

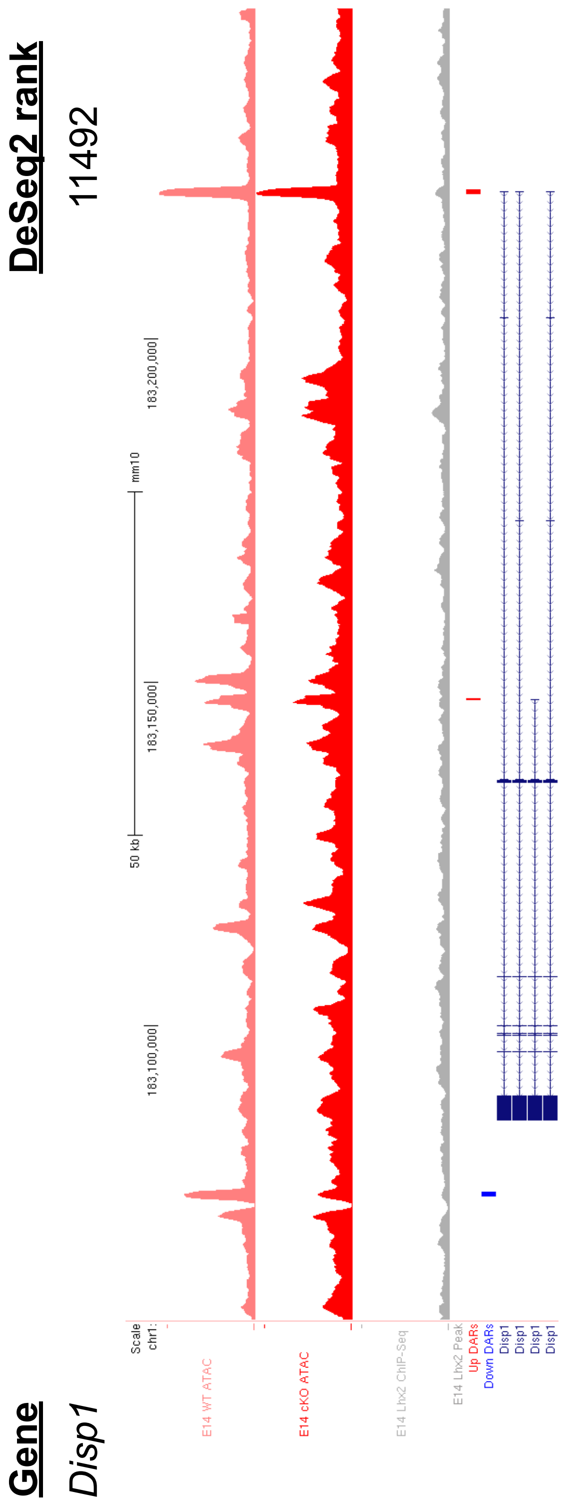

Supplemental Figure 4

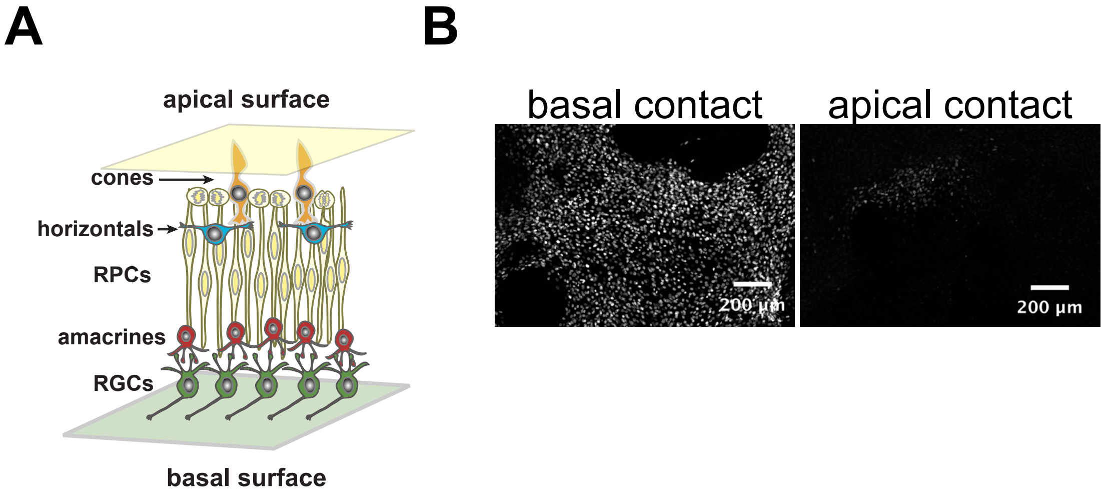

Supplemental Figure 5

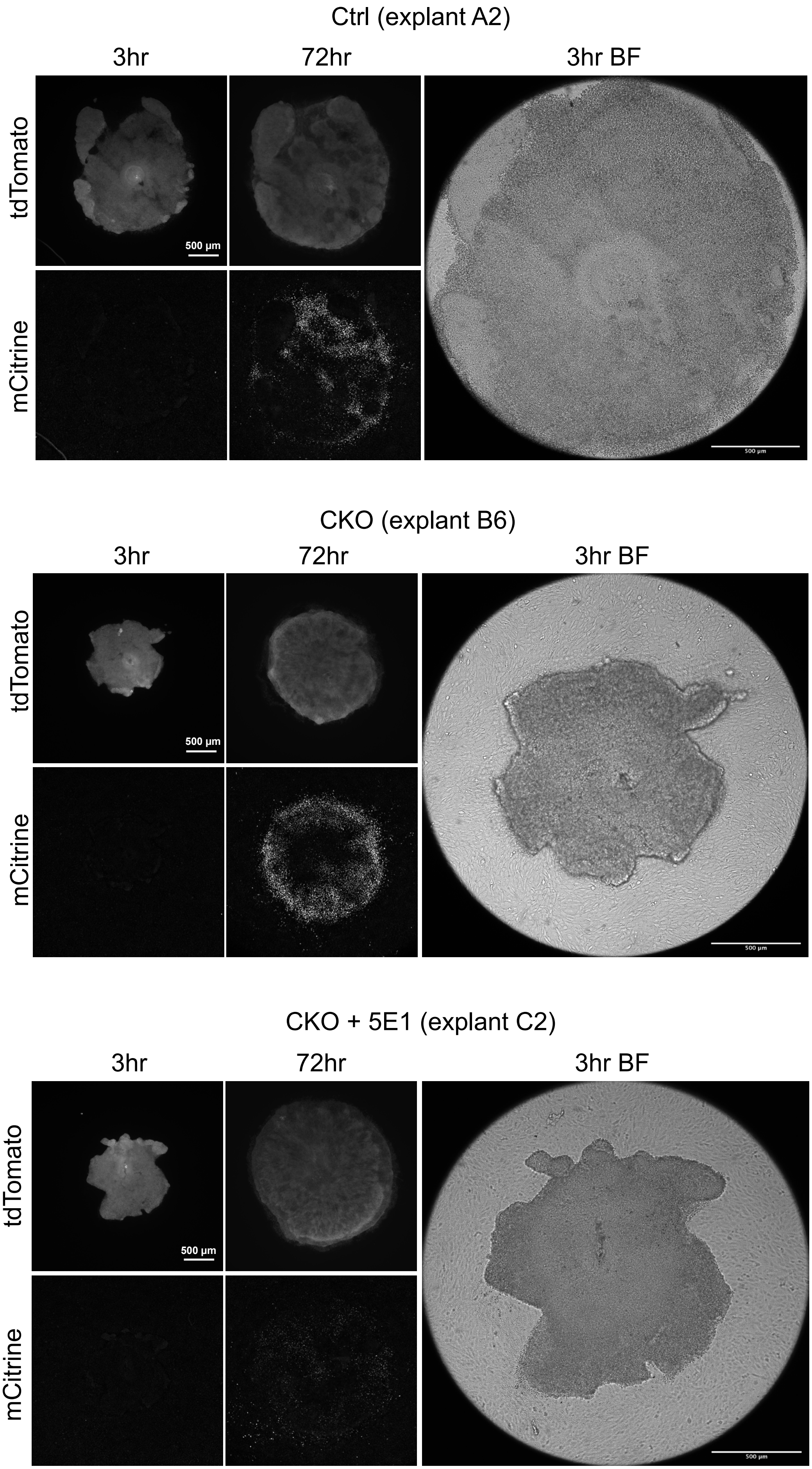

Supplemental Figure 6

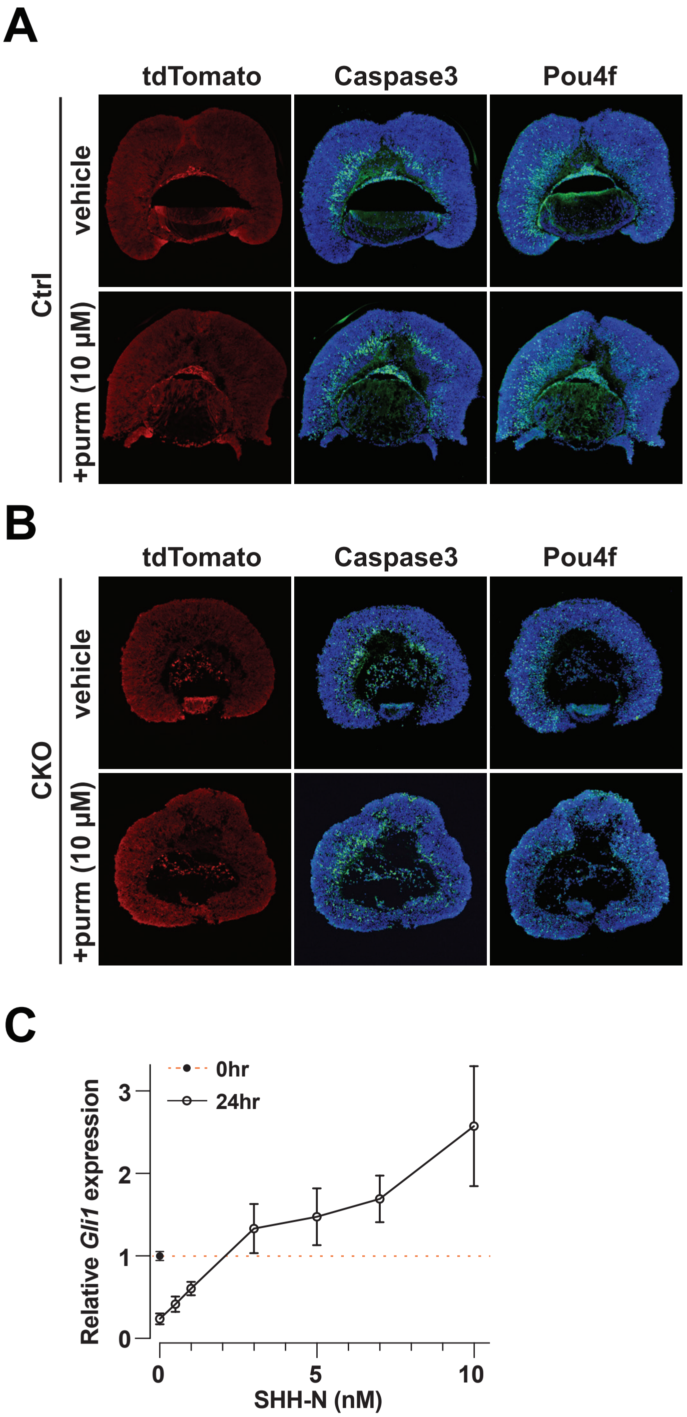

Supplemental Figure 7

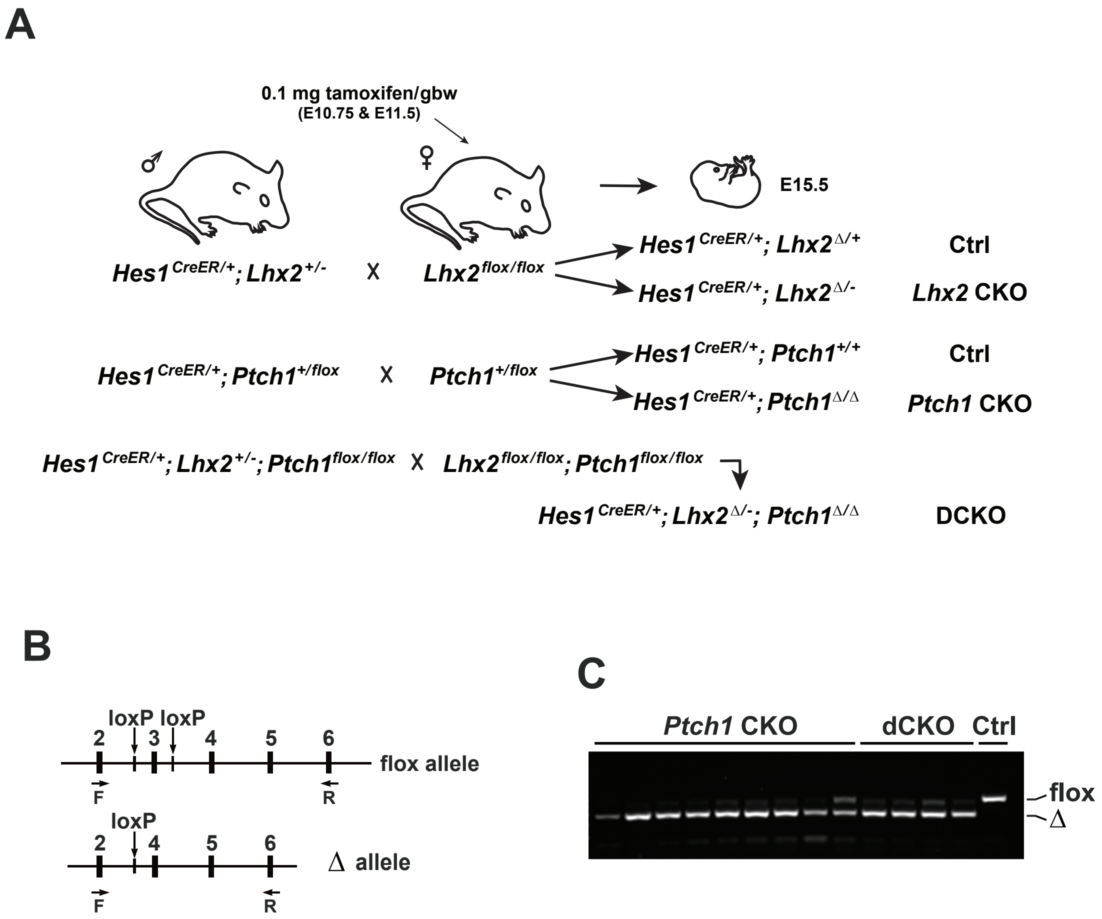

Supplemental Figure 7

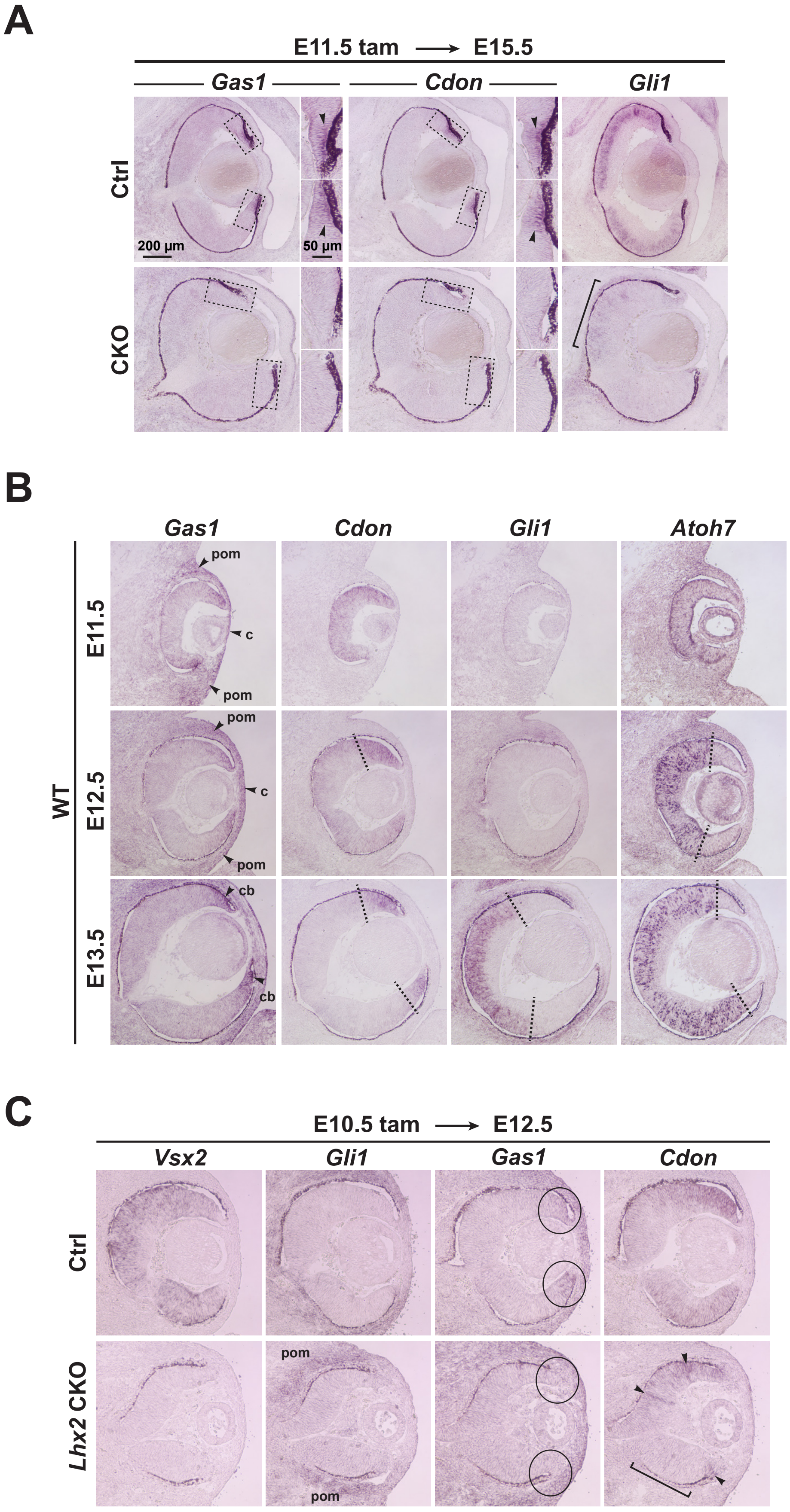
